# Chi.Bio: An open-source automated experimental platform for biological science research

**DOI:** 10.1101/796516

**Authors:** Harrison Steel, Robert Habgood, Ciarán Kelly, Antonis Papachristodoulou

## Abstract

The precise characterisation and manipulation of *in vivo* biological systems is critical to their study.^1^ However, in many experimental frameworks this is made challenging by non-static environments during cell growth,^2, 3^ as well as variability introduced by manual sampling and measurement protocols.^4^ To address these challenges we present Chi.Bio, a parallelised open-source platform that offers a new experimental paradigm in which all measurement and control actions can be applied to a bulk culture *in situ*. In addition to continuous-culturing capabilities (turbidostat functionality, heating, stirring) it incorporates tunable light outputs of varying wavelengths and spectrometry. We demonstrate its application to studies of cell growth and biofilm formation, automated *in silico* control of optogenetic systems, and readout of multiple orthogonal fluorescent proteins. By combining capabilities from many laboratory tools into a single low-cost platform, Chi.Bio facilitates novel studies in synthetic, systems, and evolutionary biology, and broadens access to cutting-edge research capabilities.

---

Biology faces a crisis of reproducibility, caused in part by a lack of control over conditions experienced by cells prior to and during experiments.^5^ Biological systems are often characterised using batch-culture methods,^6, 7^ which make isolating a cellular sub-system’s behaviour from that of its host challenging.^3, 8^ To improve the robustness of biological data an ideal experimental setup would provide a controlled, static environment in which culture parameters such as nutrient availability and temperature are regulated,^9^ and perform frequent and accurate measurements *in situ*. This can be partially achieved using continuous culture devices such as a Turbidostat, which dilutes cells during growth to maintain a constant optical density (OD). In recent years a number of Turbidostat platforms have been developed, and are beginning to find widespread applications in systems, synthetic, and evolutionary biology.^10–15^ However, in many cases these devices are not easy to build/obtain, require interfacing with external hardware (such as incubators), are inflexible for applications beyond their designed purpose, and lack *in situ* measurement and actuation capabilities (such as optogenetic actuation or measurement of fluorescent proteins) which are fundamental to many experimental studies.

To address these challenges we developed Chi.Bio, a parallelised all-in-one platform for automated characterisation and manipulation of biological systems. It is open-source and can be built from printed circuit boards (PCBs) and off-the-shelf components for ∼ $300 per device. The platform comprises three primary components (Fig. 1a); a control computer, main reactor, and pump board. The control computer can interface with up to eight reactor/pump pairs in parallel, allowing independent experiments to be run on each. It also hosts the platform’s operating system, which provides an easy-to-use web interface for real-time control and monitoring of ongoing experiments. The main reactor contains most of the platform’s measurement and actuation sub-systems (Fig. 1b), which operate on standard 30 mL flat-bottom screw-top laboratory test tubes (with a 12 to 25 mL working volume). All measurement and actuation systems (with the exception of the heat plate) are non-contact, minimising sterilisation challenges and allowing test tubes to be hot swapped during operation. Each reactor can accept up to four liquid in/outflow tubes, which are driven by peristaltic pumps housed in the reactor’s dedicated pump board. The platform as a whole is entirely modular (the three components inter-connect via micro-USB cables), allowing it to be tailored to a wide variety of experimental configurations.

**Fig. 1:**
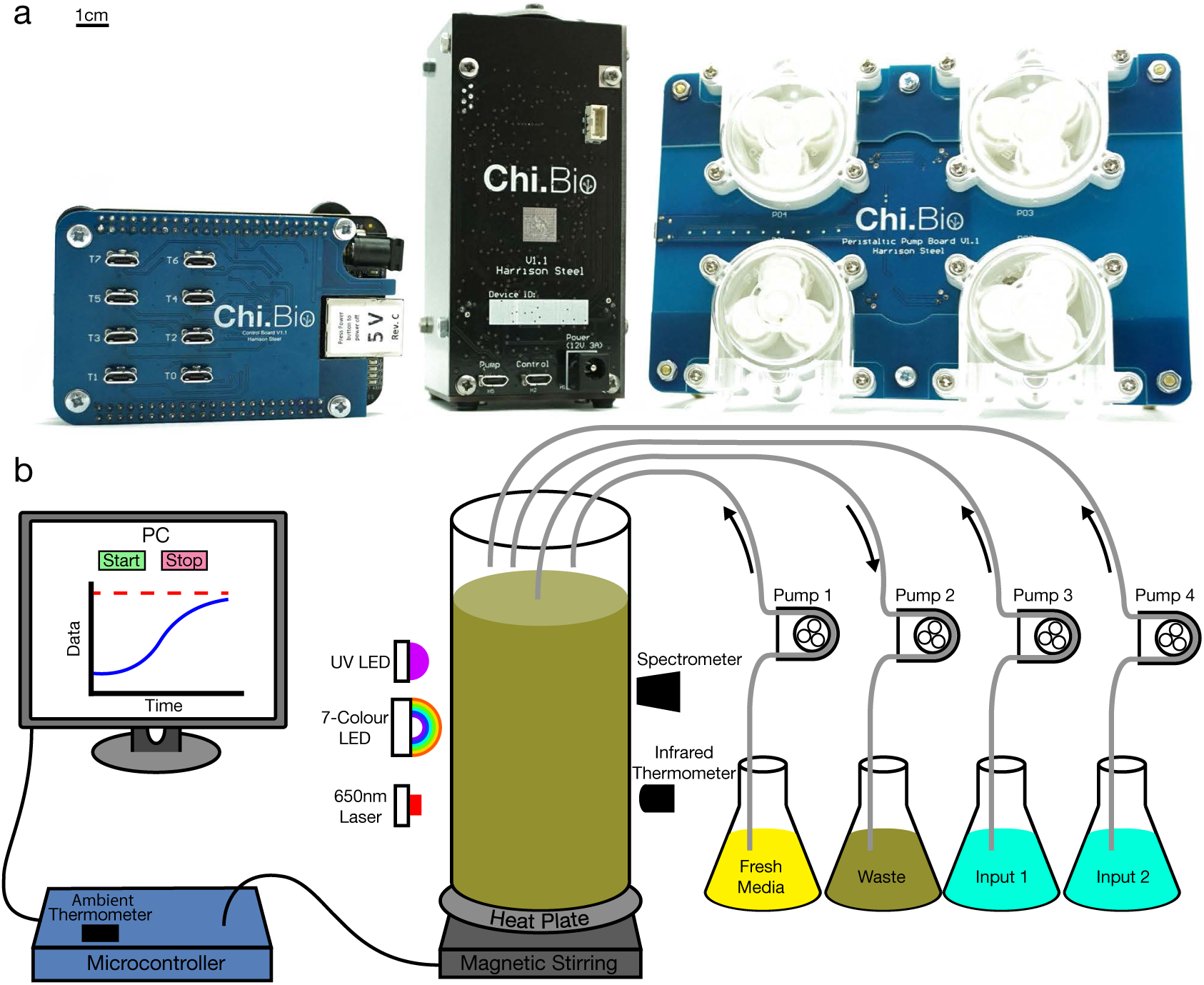
**(a)** The Chi.Bio platform comprises a control computer (left), main reactor (centre), and peristaltic pump board (right). The system is open-source and can be constructed for ∼ $300 using only PCBs and off-the-shelf components. Scale bar indicates 1 cm, giving the main reactor dimensions of 11.5×5.3×5.3 cm. **(b)** Schematic of sub-systems and interconnections. A lab computer or network connects to the control computer, which runs the platform’s operating system and can interface with up to eight reactor/pump pairs in parallel. Each reactor has a 12 to 25 mL working volume and contains a range of measurement and actuation tools for precise *in situ* manipulation of biological systems. These include a UV LED, a 650nm laser (for OD measurement), seven coloured LEDs in the visible range (for optogenetics and fluorescence measurement), and a spectrometer. An infrared thermometer and heat plate are used to regulate temperature, and the culture is agitated using magnetic stirring. Each reactor has a modular pump board with four direction- and speed-controllable peristaltic pumps. For a detailed descriptions of each hardware sub-system see Notes S1-S4.

Experimental techniques which exploit interactions between light and life, such as light-sensitive proteins, optogenetics, and fluorescent reporters, are ubiquitous in biological research.^16, 17^ Chi.Bio contains an array of optical outputs and sensors to support these techniques (Fig. 2a). A 650nm laser is used for optical density measurement (calibrated against a Spectrophotometer, Note S5), and is driven by an analogue feedback circuit (Fig. S6) to provide stable, temperature-insensitive readings (Note S6). Optogenetic actuation and excitation of fluorescent proteins employs a focused high-power 7-Colour LED with six emission bands across the visible range and a 6500K white output (Fig. 2b,c), and a separate 280nm UV LED is included to stress/sterilise cells. Each LED has an independent current-limiting pulse width modulation (PWM) driver (Fig. S6), allowing its intensity to be regulated over three orders of magnitude (Fig. S7), and is thermally coupled to the outside of the device to reject heat when operating at high intensity. The combined LED implementation provides optical outputs that exhibit minimal power and spectral variability between devices or environmental conditions (Note S7).

**Fig. 2:**
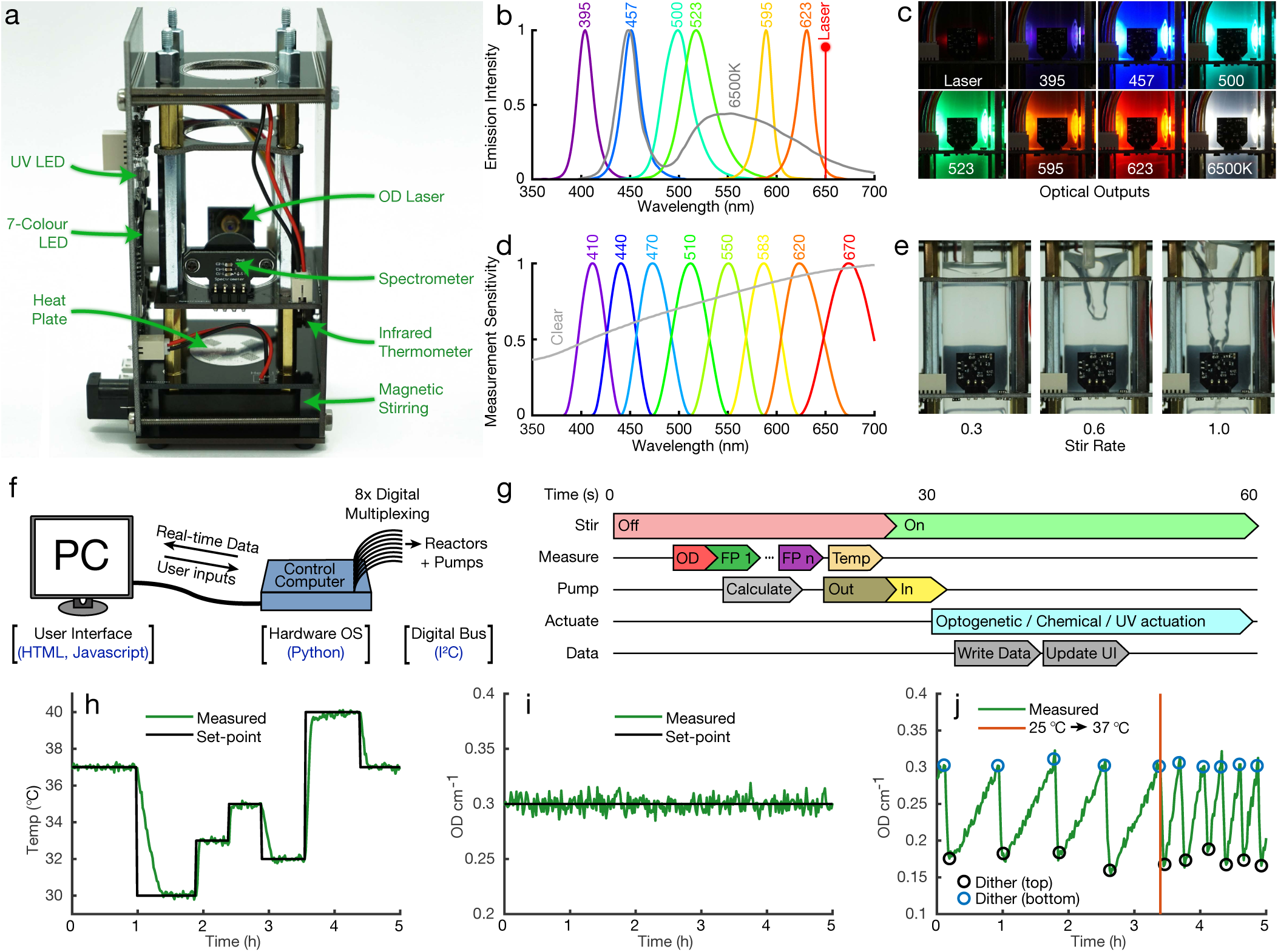
Hardware sub-systems, software, and automation. **(a)** The main reactor with sides removed and sub-systems labelled. The vertical PCB on the left hosts driving circuitry for many sub-systems, as well as power regulation and filtering. **(b)** Emission spectra of the optical outputs in the device (280nm UV LED not shown). **(c)** Images of optical outputs, with laser set to 50% and LEDs to 5% intensity. LEDs are focused and perpendicular to the spectrometer to maximise faint fluorescence signals. **(d)** Measurement filter bands of the platform’s spectrometer. **(e)** Magnetic stirring provides a powerful vortex when set to a high rate. **(f)** Software architecture, which packages multiplexed low-level commands (digital communications following I^2^C standard) into an easy-to-use web interface accessed from a connected PC or network. **(g)** A typical 60 second experimental automation cycle. Initially stirring is disabled so liquid can settle, reducing noise in measurements and providing a flat surface for removal of waste media. **(h)** Media temperature controlled to follow a pre-defined path over 5 hours. Heat input is provided by the heat-plate, cooling is passive. **(i)** The OD of *E. coli* in exponential growth phase, maintained within ∼ 2% of its set-point. **(j)** OD can be set to follow a dithered waveform (with cells rapidly diluted to a lower OD value whenever the upper limit is reached) to accurately measure growth.^18^ Following a change in temperature set-point from 25 °C to 37 °C growth accelerates significantly.

Measurements of light intensity are performed within the device by a chip-based spectrometer with eight optical filters covering the visual range, as well as an un-filtered “Clear” sensor (Fig. 2d). Multiple wavelength bands can be measured simultaneously, each with electronically adjustable gain and integration time. The spectrometer is set up to perform temperature and long-term baseline calibration using a dark photodiode prior to every measurement. Typical spectrometer measurements (e.g. of fluorescence) are reported as the ratio of light intensity measured at the fluorescent protein’s emission band to the total intensity of the excitation source; this ratiometric measurement mitigates the impact of differing excitation intensity between devices and spatial variations in culture density.

Culture temperature is measured non-invasively by a medical-grade infrared thermometer, which is accurate to ± 0.2 °C for temperatures near 37 °C. There are also Air temperature thermometers (± 0.5 °C accuracy) within the main reactor and on the control computer for monitoring the surrounding environment. Temperature change is actuated by a PCB-based heat plate, capable of heating a 20 mL culture at up to 2.0 °C min^−1^. Below the heat plate is a magnetic stirring assembly built upon an off-the-shelf fan, which has an adjustable stirring rate (Fig. 2e) and can be used with standard laboratory stir bars. The main reactor also includes an external expansion port, which provides power and a digital interface for user-built add-ons to Chi.Bio.

External to the main reactor is a pump board, which can house up to four low-cost peristaltic pumps with independently controllable speed (up to 1 mL s^−1^) and direction. Each pump transfers liquid to/from the culture test tube via standard 4.5mm silicone laboratory tubing, which can be installed without joints from input to output to assist sterilisation. Typically two pumps are dedicated to turbidostat functionality (one for input of fresh media, one for removal of waste), leaving the other two free for programmable mixing of media/inducers, or transfer of liquid between reactors. Detailed specifications of Chi.Bio’s hardware and electrical sub-systems are outlined in Notes S1-S4, and analysis of their calibration, measurement stability, and inter-device variability is described in Notes S5-S7.

At the lowest level of Chi.Bio’s open-source operating system is a multiplexed I^2^C bus for digital communications within the device (Fig. 2f). Digital signals are in turn controlled by the hardware operating system (Python, Note S8), which implements automation functions as well as data collection, processing, and storage (in .csv format). Real-time data is output periodically via a web-server (accessible from a connected PC or network) which provides an easy-to-use web user interface (built in HTML/Javascript, Note S9). The complete software stack lets users set up, control, and monitor standard experiments entirely through their web browser, and more advanced protocols or algorithms which combine any of the platform’s measurement/actuation capabilities can be easily implemented using the in-built “custom program” Python framework (Note S10).

A typical automated experiment follows a 60 second cycle (Fig. 2g) during which the stirring vortex is allowed to settle prior to measurement, media/waste is added/removed, and data is processed to calculate new control inputs. This automation protocol can be adjusted in real-time to change controller set-points or measurement setup, all of which are recorded throughout an experiment. Automated temperature and OD regulation are implemented using PID and Model Predictive Control (MPC) algorithms.^20^ The temperature control algorithm (Note S11) facilitates rapid temperature changes with minimal error and set-point overshoot (Fig. 2h). The OD control algorithm (Note S12) typically maintains OD within ∼2% of its set-point (Fig. 2i), and can also be used to dither a culture’s OD about a central set-point (Fig. 2j) for precise measurement of growth rate at near-constant density.^18^

A straightforward application of Chi.Bio is the study and regulation of growth in changing conditions. This can involve the collection of growth curves (Fig. 3a), or dithering of OD near a set-point (as in Fig. 2j) to analyse the dependence of growth on a particular parameter (e.g. temperature, Fig. 3b). The UV output can be used to stress cells (or encourage mutagenesis^21^); in Fig. 3c *E. coli* is subjected to different UV intensities, causing a gradual reduction in growth. Chi.Bio can also be used to monitor the growth of biofilms (using the procedure outlined in Note S14); in Fig. 3d we observe that initial adaptation to the continuous culture environment (which actively selects for biofilm-forming phenotypes^22^) leads to accelerated biofilm formation.

**Fig. 3:**
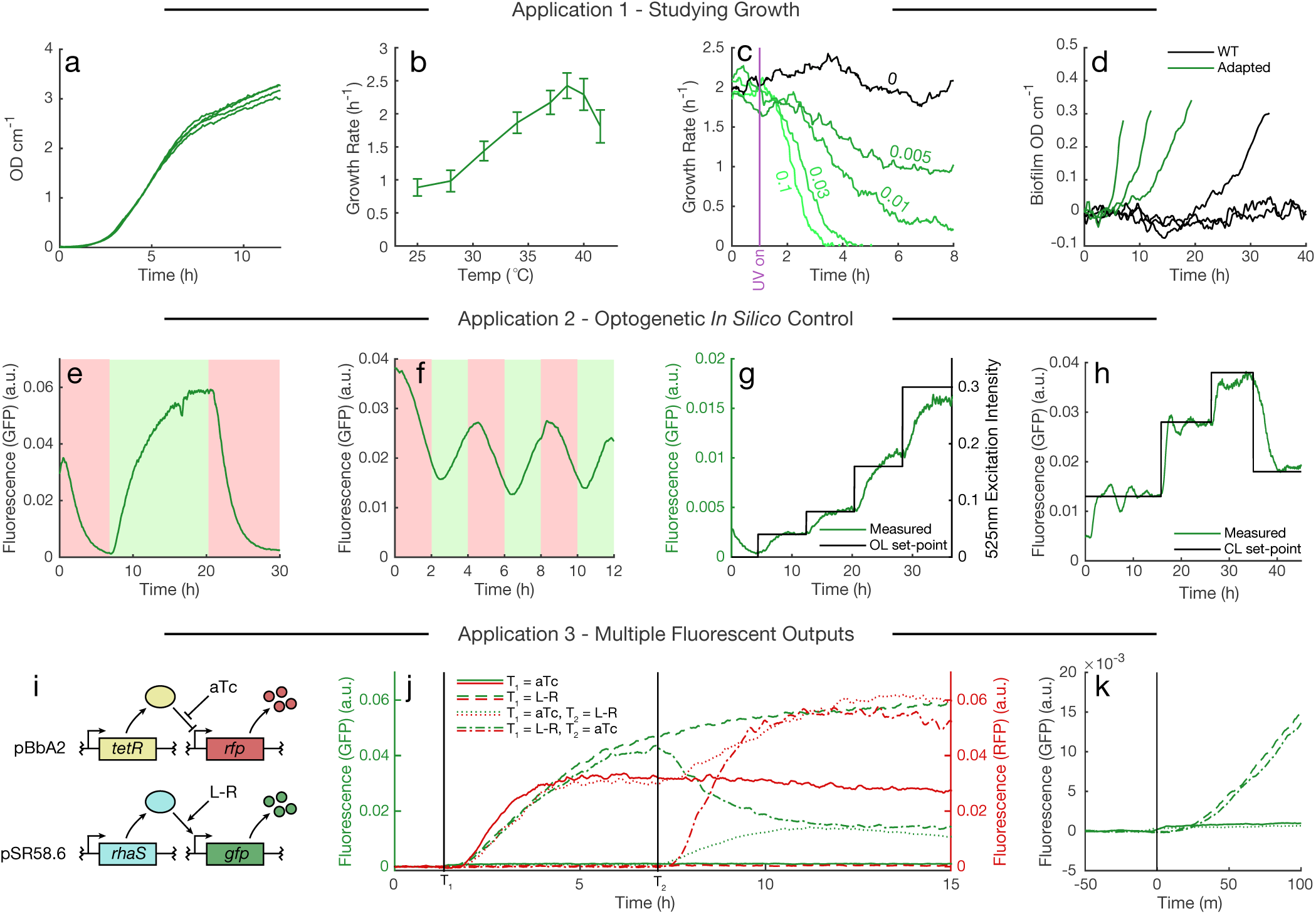
**Application 1 - (a)** Growth curves of *E. coli* (four replicates). **(b)** Dependence of *E. coli* growth rate on temperature, error bars represent standard deviation of growth rates measured over five hours at each temperature. **(c)** Measured growth rate following activation of UV source at specified power level at *t* = 1 hour. The population is able to adapt to low UV intensities and eventually returns to its initial growth rate (Fig. S23). **(d)** Biofilm optical density versus time for cells before and after adaptation in Chemostat mode for 120 hours, calculated as described in Note S14. **Application 2 - (e**,**f)** Optogenetic CcaS-CcaR system coupled to GFP expression,^19^ stimulated with slow- and fast-varying inputs (green/red light activate/deactivate gene expression respectively). **(g)** Fluorescence expression (left axis) controlled by varying optogenetic excitation intensity (right axis) in open loop (OL). **(h)** Fluorescence expression regulated in closed loop (CL) to follow a pre-determined profile, using a proportional-integral (PI) controller. **Application 3 - (i)** Two-plasmid system for inducible expression of GFP and RFP. **(j)** Fluorescence of GFP (left axis) and RFP (right axis) following induction at times T1 and T2 with indicated inducer combinations. **(k)** GFP fluorescence measured during short time period near *T*1. A small increase is observed in the two samples to which aTc is added due to the fluorescence of the inducer compound itself.

Real-time interfacing between *in silico* computational elements and biological systems can facilitate novel studies^23^ and optimise experimental schemes.^24^ We implemented such a scheme as a “custom program” in the platform, which uses the red (623nm) and green (523nm) LEDs as actuating inputs for the CcaS-CcaR optogenetic system coupled to GFP expression.^19^ In Figs. 3e,f this system is probed with square wave inputs of varying frequency, which yield smoothed output signals (due to the dynamics of protein expression/maturation). In Fig. 3g an input of varying intensity is supplied, revealing a non-linear dependence between fold-change of input versus output. This highlights a major challenge posed by open-loop control of biology; predicting *a priori* the behaviour of such complex, non-linear systems requires accurate and implementation-specific models,^6, 25^ which may only be relevant in a limited range of tightly controlled environmental conditions. This can be overcome by *in silico* feedback control:^12^ In Fig. 3h our custom program implements a control law which updates the optogenetic excitation intensity each minute depending on the measured fluorescence, steering the culture’s fluorescence to follow a complex profile without requiring a model or extensive *a priori* analysis of the biological system itself.

Characterisation of biological systems often requires monitoring of multiple fluorescent protein-tagged outputs.^7, 26, 27^ Here we demonstrate the effectiveness of Chi.Bio for probing such a system (Fig. 3i). Adding chemical inducers in different orders highlights the impact that cellular burden^7^ has on its two outputs (Fig. 3j): if RFP is induced after GFP we observe a significant drop in GFP (due to limited resource availability for production of GFP and its activating transcription factor RhaS). However, if RFP is induced first the subsequent induction of GFP leads to an *increase* in RFP expression, which we hypothesise is due to the effect of resource limitation on expression of its repressing transcription factor TetR (repeatability of this behaviour is illustrated in Fig. S28). In all cases we observe negligible cross-talk between fluorescent protein measurement channels, and our instrument’s high sensitivity allows fluorescence activation to be observed within ∼ 20 minutes of induction (Fig. 3k, approximately the time required for fluorophore maturation).

The applications outlined herein represent only a fraction of Chi.Bio’s potential use-cases. The platform’s turbidostat functionality lends its application to continuous directed evolution^28^ (as demonstrated by eVOLVER^13^) which could be tuned using the UV output.^21^ Direct integration of fluorescence measurement with chemical/optogenetic actuation and computational capabilities facilitates optimised experimental design^25^ and the implementation of online experimental-planning algorithms.^24^ Given that the only components of Chi.Bio which come into direct contact with cells are standard laboratory consumables (glass test tubes, silicone tubing, magnetic stir bars), such experiments could be run with a broad range of cell types and reagents (e.g. algae, with photosynthesis supported by the tunable LED outputs). A list of potential applications (and minor hardware modifications that can be made to enable others) is presented in Table S1.

The cost and resulting inaccessibility of modern scientific hardware is one of the primary obstacles to participation in cutting-edge research worldwide, particularly in small laboratories and institutions or developing nations.^29^ Consequently we have made Chi.Bio entirely open-source, with schematics, code repositories, user manuals, and a public support and discussion forum all available on the project’s website (https://chi.bio). A single Chi.Bio device (consisting of one control computer, one reactor, and one pump board) can be assembled by hand for ∼ 300, or can be purchased ready-built from a supplier of open-source scientific hardware. In the long term we hope that Chi.Bio will provide a versatile tool for biological sciences, and broaden access to cutting-edge research capabilities.

## ACKNOWLEDGEMENTS

The authors would like to thank Prof Wei Huang for use of his laboratory space.

## AUTHOR CONTRIBUTIONS

H.S. conceived the study and performed all hardware and software engineering. R.H., H.S., and C.K. planned and conducted biological characterisation experiments. All authors wrote the manuscript.

## METHODS

### Strain and plasmid selection

*Escherichia coli* strain MG1655 was used for growth/biofilm characterisation experiments. Fluorescent protein experiments utilised BL21(DE3) *E. coli*, apart from the optogenetics experiments which utilised BW29655. Bacterial strains were made chemically competent by treatment with calcium chloride and transformations were performed via heat shock. All *E. coli* strains used in this study are freely available (Table S2), as are plasmids (Table S3).

### Culture techniques

Cells were cultured in either LB (growth/biofilm experiments), or EZ rich defined media (Teknova Inc; cat: M2105) supplemented with 1% (v/v) glycerol (for optogenetic/fluorescence experiments). All experiments were performed in 20 mL volume in 30 mL test tubes (Fisher Scientific, 11593532) open to atmosphere. Stirring employed disk stir bars (Fisher Scientific, 11878892) at speed setting 0.6. Culture temperature was maintained at 37°C unless otherwise stated. For fluorescent protein and optogenetic experiments cells were maintained at an OD of 0.4. Carbenicillin (a semi-synthetic analogue of ampicillin with greater stability), chloramphenicol, and spectinomycin were used at final concentrations of 100 *µ*g mL^−1^, 25 *µ*g mL^−1^, and 50 *µ*g mL^−1^, respectively.

### Experiment setup procedure

Prior to experiments test tubes and stir bars were sterilized by autoclave. Silicone tubing was sterilised by pumping 70% ethanol for 10 seconds, and media for 5 seconds. For experiments that required chemical induction (Application 3) an additional 500 mL of tap water was pumped through each tube prior to the experiment to prevent inducer cross-contamination. Tubing exterior was sterilised via swabbing with 70% ethanol. For each media type, each reactor’s OD zero point was calibrated with a test tube of fresh media as described in Note S5. Test tubes were filled with 20 mL media and inoculated prior to insertion into each reactor.

### Biofilm experiments

In each trial cells were initially diluted to an OD of 0.1, and (once reached) maintained at an OD of 0.5. Each experiment’s zero-time is defined as the time when the culture first reached OD 0.5 (typically ∼ 2 hours post-inoculation). To develop a biofilm forming phenotype *E. coli* was grown at high density (OD > 1) in LB media in a reactor in chemostat mode for 120 hours. Samples from this culture were subsequently used to inoculate each of the “adapted” biofilm trials.

### Fluorescence measurements

GFP was excited by the 457nm LED at power setting 0.1, and measured at x512 gain and 0.7s integration time using the Clear filter as base-band and 550nm filter as emission band. RFP was excited by the 595nm LED at power setting 0.1, and measured at x512 gain and 0.7s integration time using the Clear filter as base-band and 670nm filter as emission band. A baseline fluorescence (corresponding to the fluorescence measured for wild-type cells without plasmid) was subtracted from each measurement. Measurements were then smoothed using a moving mean filter with 10 minute width. Examples of raw (unprocessed) data are presented in Fig. S27. For a detailed analysis of the fluorescence measurement procedure see Note S15.

### Optogenetic control

In optogenetic experiments which required full activation/deactivation (Fig. 3e,f) either the green (525nm) or red (625nm) LED was enabled individually at power setting 0.1. In experiments where the precise induction level was controlled (Fig. 3g,h) the procedure suggested by Olson *et al.*^6^ was followed; the red LED was illuminated at a fixed power setting of 0.1, and the green LED’s intensity was varied.

### Chemically induced reporters

In fluorescence experiments which used chemically-inducible fluorescent reporters L-Rhamnose concentrations of 0.1 mg mL^−1^ and aTc concentrations of 300 nM were used. At the time of induction each inducer was added manually to both the reaction chamber and the fresh media supply (to provide a sharp step-increase) without stopping the experiment.

