## Supplementary Material for "Chi.Bio: An open-source automated experimental platform for biological science research"

### Chi.Bio

**Supplementary References** **43**

### Overview

This document contains supplementary figures and data for the Chi.Bio experimental automation platform (Fig. S1). Additional documentation and manuals for the Chi.Bio platform are available on the official website, http://chi.bio. In particular, documents of interest include:

- Getting Started Guide (https://chi.bio/getting-started/) - Overview of approaches to getting Chi.Bio set up and running, which can be done either via self-assembly or purchase of a pre-built device.

- Hardware Build & Assembly Manuals (https://chi.bio/hardware/) - Schematics of the platform, as well as manuals for manufacturing, assembly, and mechanical setup.

- Software Setup Manual (https://chi.bio/software/) - Manual for setup of the operating system and software, including links to the Github repository for latest versions.

- Operation Manual (https://chi.bio/operation/) - Manual for experimental operation of the platform, including notes on software functionality, sterilisation procedures, calibration, and data processing.

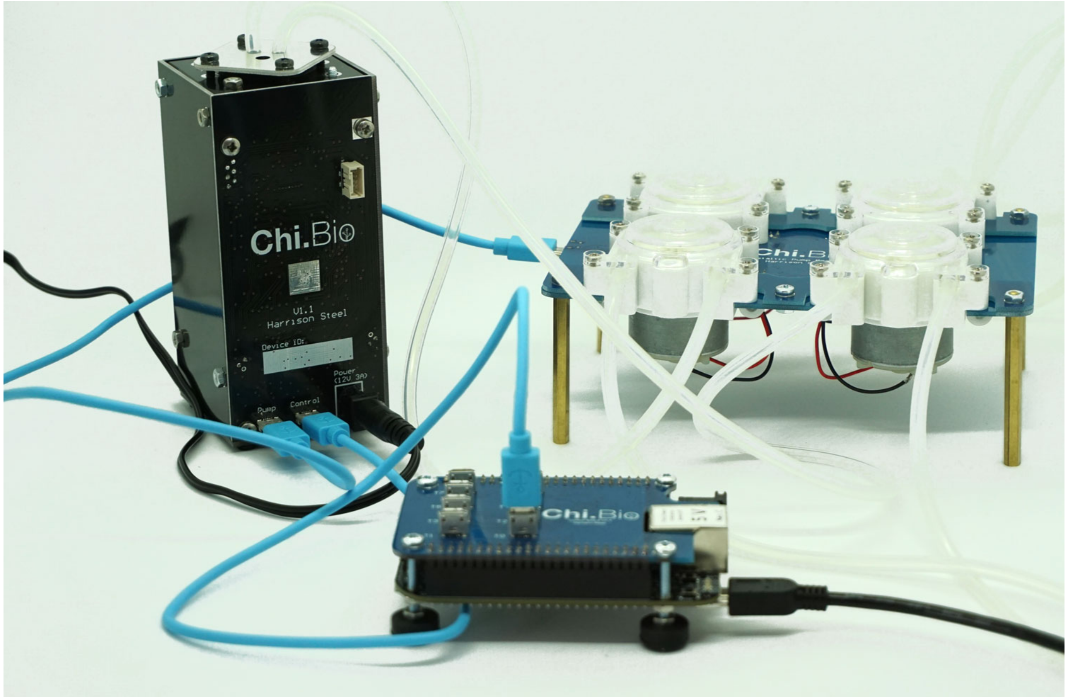

Figure S1: **Chi.Bio.** Chi.Bio setup with all connections/tubing. Components interface with one another via microUSB cables, and power is supplied to the main reactor by a 12V wall adapter.

### Hardware

#### Note S1 Electrical Architecture

Chi.Bio contains many interlinked electrical sub-systems, which are spread across nine printed circuit boards (PCBs) and a Beaglebone Black microcontroller. The modularity of the main components (Control Computer, Main Reactor, Pump Board) allows the platform to be re-configured and scaled to suit a range of experimental needs. The electrical connections between each sub-system, as well as the contents of each PCB, are summarised in Fig. S2. Many of these PCBs have secondary purposes, such as providing structure or positioning for components within the device.

Power is supplied to the platform by an off-the-shelf 12V power adapter (with short-circuit protection), which is down-regulated to provide a number of low-noise power rails for individual components (discussed in Note S3). Digital communications within the platform follow the $I^2C$ bus standard, which uses two 3.3V digital lines (SDA, serial data, and SCL, serial clock) for bidirectional data transfer. There is a single $I^2C$ "master" (the Beaglebone) which controls communications with each $I^2C$ "slave" (the other digital electronic sub-systems) according to the slave's address on the bus. The platform's operating system (Note S8) includes data handling and library functions to control all communications on the $I^2C$ bus, meaning the user can program new features using high-level commands only (and not worry about their low-level digital implementation).

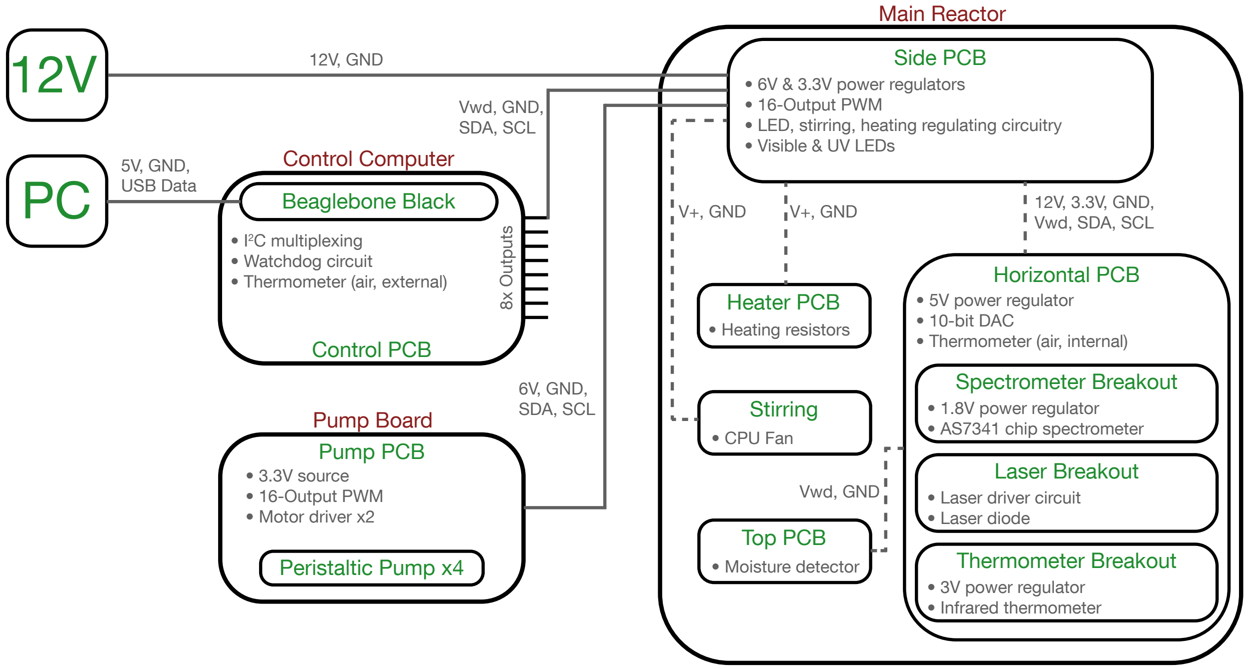

Figure S2: **Chi.Bio Hardware Architecture.** Electrical layout of the Chi.Bio system, indicating each electronic subsystem and its connectivity. The Control Computer (Note S2), Main Reactor (Note S3), and Pump Board (Note S4) assemblies are labelled in red, and each sub-system in green. Solid/dashed grey lines indicate external/internal wired connections respectively. Each wired connection is labelled with the signals it carries. SDA/SCL are the $I^2C$ data/clock lines respectively, the V+ are independently controllable voltage levels, Vwd is the watchdog circuit's indicator voltage.

#### Note S2 Control Computer

The Control Computer (Fig. S3a) is built on an off-the-shelf Beaglebone Black microcontroller (a simple single-board Linux computer) which connects to an accompanying laboratory PC via USB. The Beaglebone has a single-core ARM Cortex-A8 Processor (which we run at 1GHz), 4GB of on-board flash memory, an SD card slot, and USB/HDMI/Ethernet connectivity. Our implementation uses the open-source Debian 10 operating system (with Linux Kernel 4.19). The Beaglebone interfaces with a custom-made control PCB (Fig. S3b) which allows up to eight Chi.Bio reactor/pump pairs to be controlled in parallel via micro-USB cables, and also hosts a digital thermometer for measurement of ambient temperature.

In order for multiple pieces of identical hardware (i.e. the Chi.Bio reactors) to be connected to a single $I^2C$ "master" (the Beaglebone) their digital busses must be systematically connected to the controller only when communications are necessary. This is achieved by an 8x digital multiplexing circuit on the control board. Also implemented on the control computer is a hardware-embedded watchdog circuit, which periodically checks that the Chi.Bio operating system is running. If this check fails (e.g. if the user creates a custom algorithm with bugs that causes the operating system to fail) then a signal from the control computer disconnects the power regulation circuitry on each reactor, preventing damage. This power-disconnect circuit is also used to sense any moisture in the device (described further in Note S3).

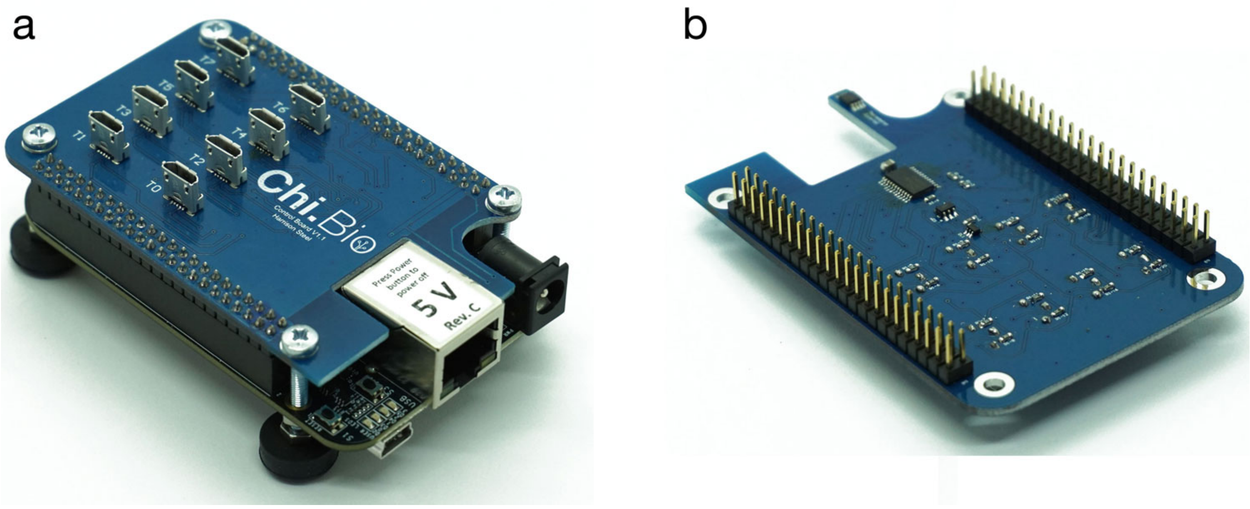

Figure S3: **Chi.Bio Control Computer.** **(a)** Complete assembly including Beaglebone Black control computer. **(b)** Underside of Control PCB which includes digital multiplexing, watchdog circuitry, and an air temperature thermometer.

### Note S3 Main Reactor

As the central component of the Chi.Bio platform, the main reactor contains many of its measurement and actuation sub-systems. It is constructed from printed circuit boards (PCBs) and off-the-shelf fasteners (nuts, bolts), allowing it to be quickly dis/assembled as required. It accepts standard 30ml flat-bottom test tubes, which can be fitted with the platform's custom-made lid or any number of commercial lid assemblies (that fit the common 22-400 thread standard). It has four external connection points; two micro-USB ports (to connect to the control computer and pump board), a 12V power supply input, and a Mini-CT expansion header (for future expansions of sensors/actuation hardware). The reactor's power regulation circuitry and drivers for the LEDs, heater, and stirring motor, as well as the LEDs themselves are implemented on its side PCB (Fig. S4b). This PCB interfaces internally (again via Mini-CT headers/cables) with the reactor's central stack (Fig. S4c) as pictured in Fig. 2a of the main text. Each reactor has a hardware-encoded digital identity number (which is bound to its infrared thermometer), allowing each reactor (and any calibration settings) to be tracked over time even as devices are physically re-shuffled. We now describe each of the reactor's sub-systems in detail.

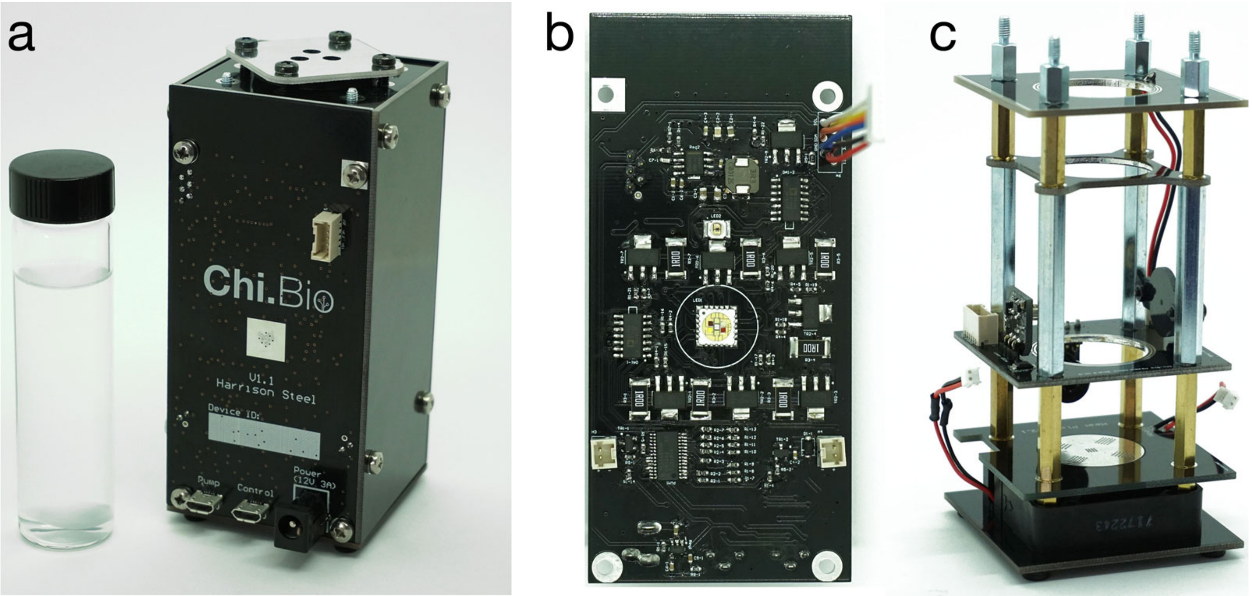

Figure S4: **Chi.Bio Reactor.** **(a)** Complete reactor with 30ml reaction tube. **(b)** Side PCB (without LED lens), which hosts much of the power and control circuitry. **(c)** Central stack with sides removed.

**Stirring:** The magnetic stirring assembly (Fig. S5a) employs an off-the-shelf computer fan, with two neodymium magnets attached. When the fan is powered this assembly generates a rotating magnetic field at the bottom of the culture flask, which rotates a magnetic stir bar in the test tube to provide aeration. The stirring fan is connected to the 6V power rail through an H-bridge motor driver, which is in turn controlled by a PWM signal that is varied to control the fan's rotation rate.

**Heating:** The reaction tube is heated using a custom-designed PCB hot plate (Fig. S5b,c). Heat is generated by passing current through four 100 Ω resistors arranged in parallel at the bottom of this PCB. There is an exposed copper pad on the upper side of the PCB, upon which the culture tube sits. Heat is transferred between the top and bottom sides of the PCBs using a large number of Vias (the copper plated holes which make electrical connection between opposite sides of a PCB) which are connected to the ground side of each resistor. Since the thermal conductivity of copper is much greater than that of fibreglass (the structural material of the PCB) most of the heat conducted away from the resistors is transferred to the upper surface of the heat plate. The heating element is connected to the 12V voltage rail in series with a MOSFET transistor, which is in turn switched by a PWM signal to vary the rate of heat production. The heat plate's maximum power output is approximately 3W, which enables rapid temperature changes of up to $\approx 2.0$ °C min$^{-1}$ for a typical 20 ml culture (Fig. 2h).

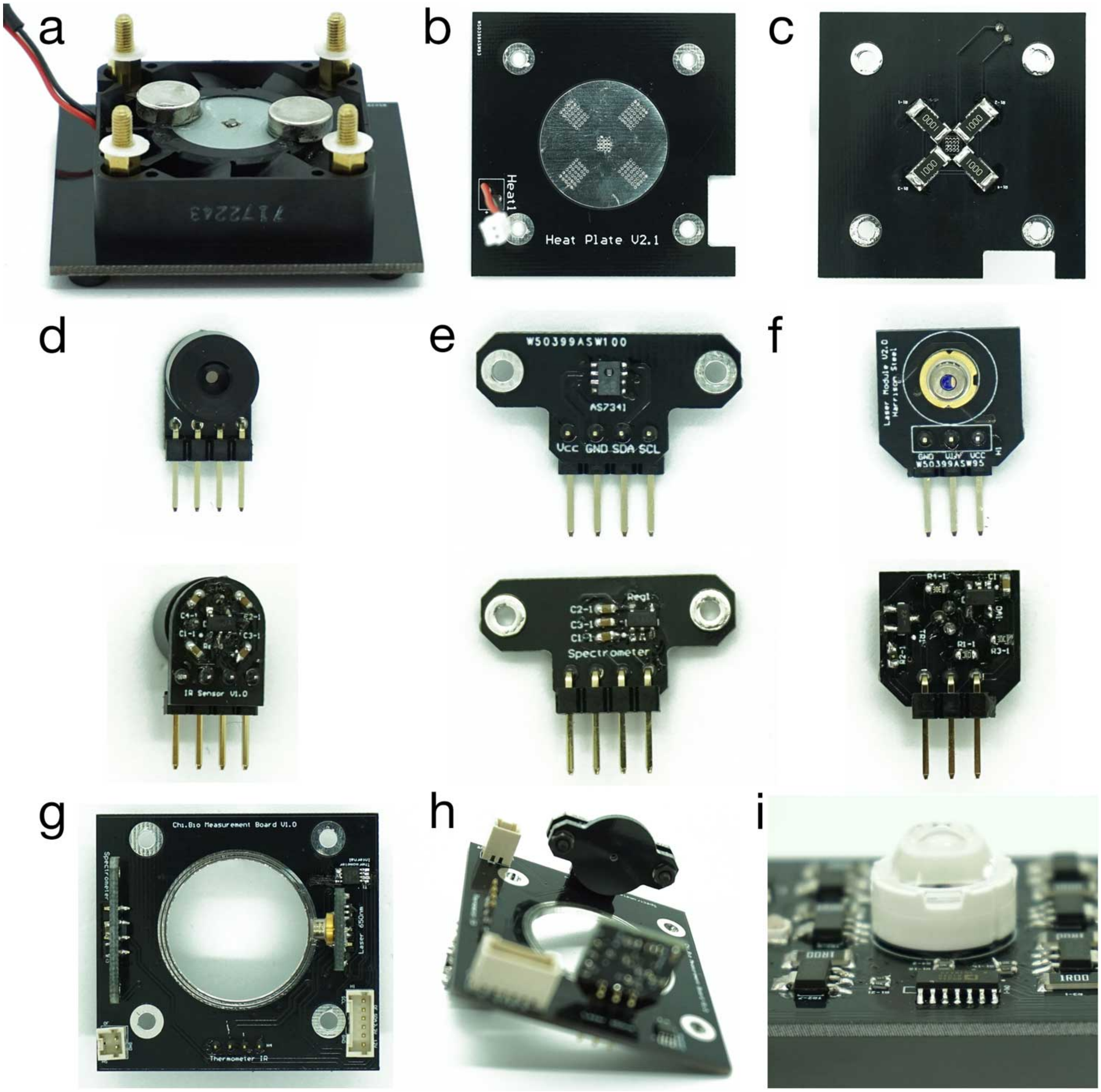

Figure S5: **Chi.Bio Reactor Subsystems.** **(a)** Stirring motor with magnets, attached to the base of the reactor. **(b)** Top and **(c)** bottom of the heat plate. **(d)** Infrared thermometer breakout board including sensor and voltage regulator/filter. **(e)** Spectrometer breakout board (without mechanical aperture) including sensor and voltage regulator/filter. **(f)** OD Laser breakout board including power control circuit. **(g)** Horizontal PCB which mounts sensors/laser, as well as the air temperature thermometer and DAC. **(h)** Horizontal PCB, now with spectrometer aperture attached. **(i)** View of side PCB (as in Fig. S4b) with visible LED Lens attached.

**Thermometry:** The temperature of the cell culture is measured directly using a medical-grade infrared thermometer (Fig. S5d) developed by Melexis.[1] We provide this thermometer with a dedicated low-noise 3V source, which allows it to remotely measure culture temperature with an accuracy of $\pm0.2$ °C for $T \in [36, 38]$ °C, $\pm0.3$ °C for $T \in [22, 36] \cup [38, 40]$ °C, and $\pm0.5$ °C outside of these ranges. The thermometer includes built-in thermal calibration (such that changes in ambient temperature do not affect its reading), and its restricted field of view (and direct proximity to the test tube) ensures that the measured temperature reflects that of the media (and not its surroundings).

**Spectrometry:** Measurement of light intensity at various wavelengths is performed by an AS7341 chip spectrometer, developed by AMS[2] (Fig. S5e). This device incorporates eight photodiodes with varying optical filters in the visible range, as well as a clear (no-filter) photodiode (summarised in Fig. 2d). The peak wavelengths/FWHM (full width half maximum) of the eight individual filters are (in nm) 410/29, 440/33, 470/36, 510/40, 550/42, 583/44, 620/53, 670/60. The spectrometer has an electronically adjustable gain (in multiples of 2) from 0.5x to 512x, and integration time (typically set to 0.7s, but can be set as low as 2.78 $\mu$s). Typically we employ a gain of 1x for OD (laser) measurement and 512x for fluorescent protein measurement, both of which return measurements in the mid-range of the spectrometer's 16 bit output. The spectrometer is set up to use an internal photodiode with a dark (light-blocking) filter to calibrate itself against temperature and long-term baseline shifts prior to each measurement. Our implementation of this sensor includes a filtered low-noise voltage source dedicated to the spectrometer, as well as a mechanically adjustable aperture (to restrict its field of view), shown attached in Fig. S5h.

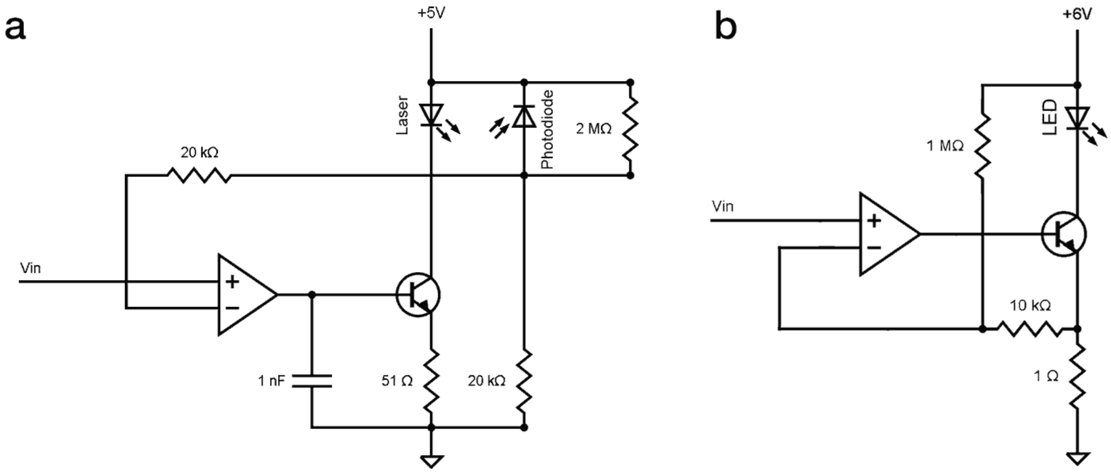

Figure S6: **Laser/LED Driver Circuits. (a)** Laser driver circuit, which uses a photodiode to maintain a constant optical power output in the presence of temperature fluctuations. **(b)** LED driver circuit, used for the 7-colour LED (and with minor adjustments for the UV emitter), which controls the current through each LED (measured across the 1 Ω resistor) to maintain a constant optical power output. A high power transistor and 1 Ω resistor are used in the LED driver circuit to handle the $\approx 0.7$ A peak current possible during operation (i.e. when the LED is set to maximum power).

**OD Laser:** We employ a de-focused 650nm laser (Fig. S5f) to measure the culture's optical density using the absorbance method (see Note S5). It is oriented antiparallel to the spectrometer, in order to provide a direct transmission measurement (as performed by a spectrophotometer or plate reader). The diode we use includes a monitor photodiode within its housing, which allows us to implement an analogue optical power regulation circuit (Fig. S6a) to reject temperature change-induced fluctuations. This circuit uses an OpAmp in feedback with a transistor to regulate the laser diode's drive current such that a fixed optical output (measured by the current generated by the laser's inbuilt photodiode) is maintained. The laser's analogue input voltage (i.e. Vin in Fig. S6a) is supplied by an $I^2C$ programmable 10-bit DAC (digital to analogue converter). We deliberately omit any focusing optics in order to generate a beam with large divergence ($> 8°$ along each axis) as this was found to result in reduced noise at the detector.

**Visible LED:** The main LED (the LZ7 emitter developed by LED Engin,[3] Fig. S4b centre) includes seven outputs in the visible range. Six of these have near-gaussian output spectra which have peak wavelengths/FWHM of (in nm) 395/30, 457/35, 496/55, 525/70, 595/25, 625/30 as summarised in Fig. 2b. There is also a 6500K white daylight LED, which can be used for growth of photosynthetic organisms.[4] Each illumination channel is independently controllable over three orders of magnitude (Fig. S7) using a current-regulating feedback circuit outlined in Fig. S6b. Each LED's input voltage (i.e. Vin in Fig. S6b) is supplied by a PWM input which is scaled down to the range [0,0.7]V, and can be run at frequencies from 200 Hz to 1.5 KHz. The LED (which is primarily designed for applications such as outdoor and stage lighting) has a large maximum output power: Our power supply and heat-removal circuitry allows each diode to be driven at up to $\approx$ 500mW optical power output (though the total optical output of all LEDs should be maintained below 1.5W for thermal reasons). That said, few applications will require such high power, and generally each LED is run at $5 \sim 10\%$ of its maximal output (e.g. for applications such as optogenetics and fluorescent protein measurement). We attach a plastic lens with a viewing angle of $\approx 20°$ to the LED (Fig. S5i) to focus its output into the spectrometer's field of view in order to improve the signal-to-noise of fluorescent protein measurements. The LED is oriented at 90° to the spectrometer, again to improve measurement signal-to-noise. Due to its lens the LED's intensity is non-uniform across the entire culture volume (i.e. it is highly concentrated at its centre, Fig. S7). However, when cells are stirred the time-averaged intensity experienced by each cell in suspension is regularised.

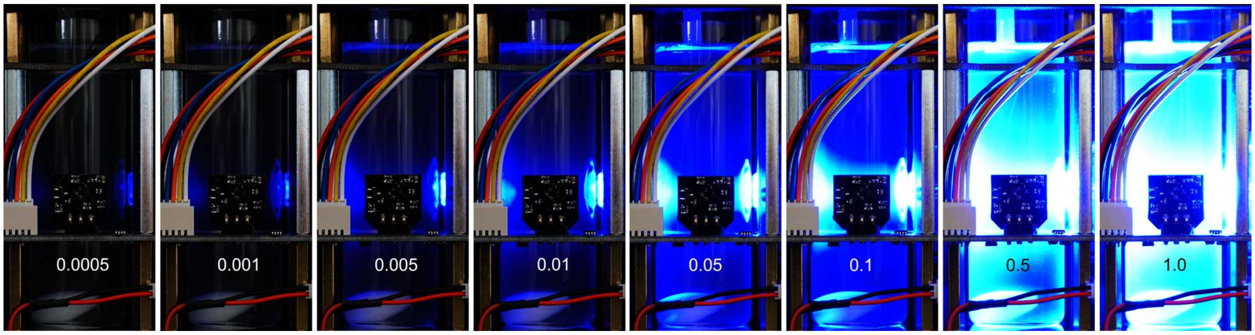

Figure S7: **LED Output vs Power Set-point.** Images of the 457nm LED (reactor sides have been removed) set to power levels between 0.0005 and 1.0, demonstrating three orders of magnitude of tunability.

**UV LED:** The UV LED (developed by RayVio[5]) emits at a peak wavelength of 280nm (FWHM 12nm), which is optimised for bacterial killing potential.[6] The emitter has a viewing angle of $\approx 120°$ and does not have any focusing optics (as we desire uniform illumination of growth media). The UV LED employs a similar current control circuit to the visible LED (Fig. S6b), with the primary difference being that the UV source is powered directly from the 12V rail such that its typical forward voltage requirement of $\approx$ 7 V can be satisfied.

**Power Regulation:** The platform's power is supplied by a 12V off-the-shelf wall adapter, which interfaces with the Side PCB of the main reactor (Fig. S4a). Inside the main reactor there are a number of different voltage regulating circuits to provide independent, low-noise supplies for many of the reactor sub-systems. In this case our design philosophy was to electronically isolate sensitive subsystems as much as possible to reduce noise/variability in their outputs/measurements. The independent power rails include the following: A 6V (max output 2A) DC-DC switching supply which is used to drive the visible LEDs, stirring fan, and pumps; a 3.3V voltage regulator which is used to drive the 16-output PWM chip on the Side PCB and send "enable" signals to all other power supplies/subsystems from the control computer's watchdog circuit; a 5V voltage regulator which supplies the OD laser only; a 3V voltage regulator which supplies the Infrared thermometer only; a 1.8V voltage regulator which supplies the spectrometer only.

**Moisture Detection:** In order to prevent moisture-related damage to the platform resulting from careless use there are moisture-detecting rings at the top and mid-point of the platform's central stack (Fig. S4c). These consist of closely spaced conductors which, if short-circuited due to contact with liquid, disable the internal 3.3V power rail (which in turn disables all other sub-systems).

### Note S4 Pump Board

The Chi.Bio peristaltic pump assembly (Fig. S8) is comprised of four peristaltic pumps with controllable speed (up to 1 mL $s^{-1}$) and direction. To achieve small rates of liquid transfer each pump is pulsed for a short period of time (minimally $\approx 0.2$ s) during each minute of operation in order to achieve a very small pump rate (when averaged over the course of that minute). The pumps transfer liquid to/from the culture test tube via standard silicone tubing with 4.5mm/2.5mm outer/inner diameter. There are no joints in the silicone tubing between its ends, facilitating effective sterilisation.

The pump board connects via micro-USB to the main reactor, which supplies it with power and digital signals (which control the pump board's inbuilt PWM driver). The pump boards are modular, allowing them to be stacked vertically as in Fig. S8c. The platform can be run in turbidostat mode without pumps 3 and 4 (which are programmable for a range of media mixing experiments), and indeed without the pump board whatsoever (if growth regulation is not desired).

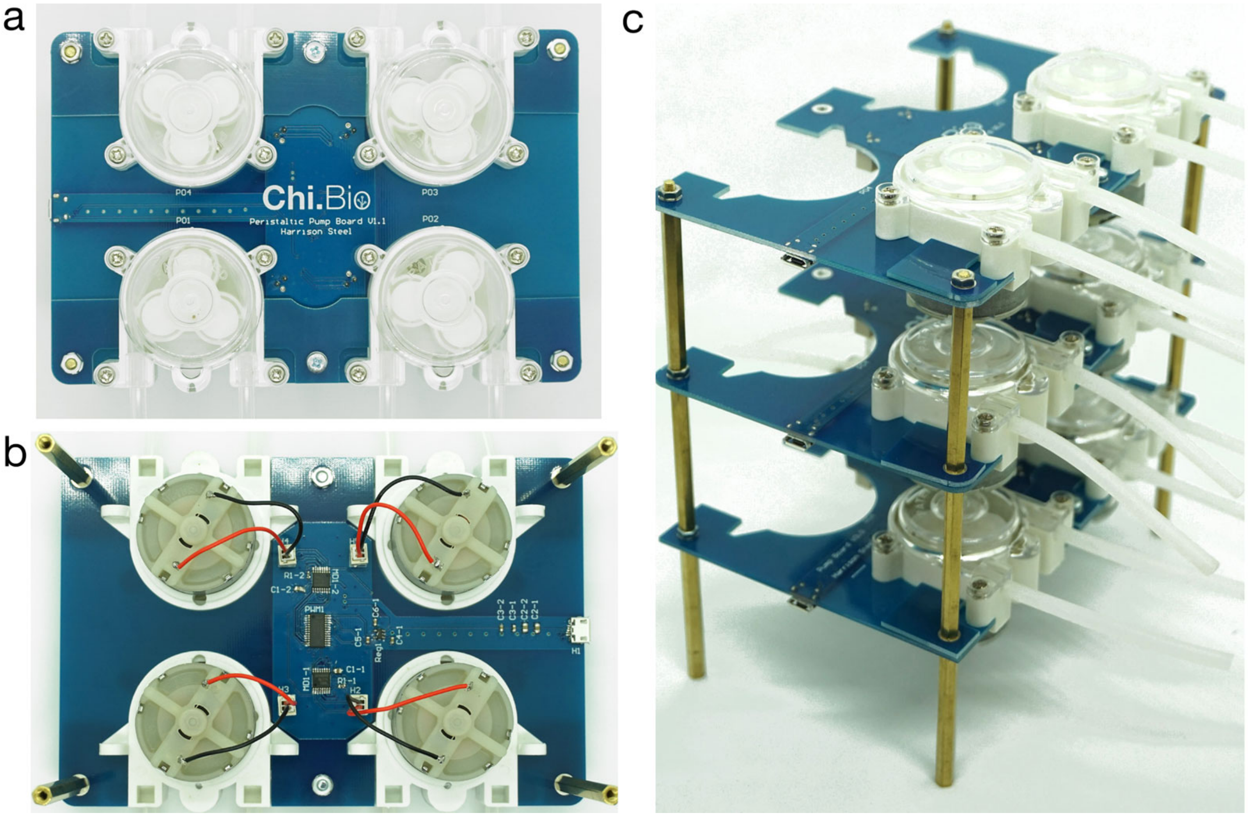

Figure S8: **Chi.Bio Peristaltic Pumps.** **(a)** Top and **(b)** bottom of the assembled peristaltic pump board, with four pumps in place. **(c)** Three pump boards (without additional pumps 3 and 4) stacked to save bench space.

### Hardware Analysis and Calibration

In this section we analyse various measurement and actuation capabilities of Chi.Bio in order to investigate anticipated levels of noise, reproducibility, and inter-device variation. We use two different calibration solutions to achieve this. The first are MacFarland standards (created by combining barium chloride and sulfuric acid in varying concentrations), which are an optical-density standard used in microbiology research.[7] One strength of MacFarland standards is their temperature stability (they change OD minimally as they are heated), however, they settle (particulates fall to the bottom of the solution) over the course of a few hours and hence require constant agitation to maintain OD. The second standard we use is evaporated milk (as suggested by Wong *et al*[8]), which has the opposite strengths/weaknesses to MacFarland standards: Milk does not tend to settle out over short time-scales, but heating can cause it to precipitate and consequently change OD over time.

#### Note S5 OD Measurement and Calibration

We calibrate the optical density (OD) readings made by Chi.Bio against a bench-top spectrophotometer (GeneQuant 1300) in order to provide standardised measurements (i.e. in units of $cm^{-1}$). There are multiple potential approaches to measuring a cell culture's OD (Fig. S9), which have been used by various experimental platforms in the past: The absorption method measures OD as the absorbed fraction of light passing through a sample (with the light source oriented antiparallel to the detector), which is the technique used in a spectrophotometer or plate reader (and used by a number of custom-built experimental platforms[9–11]). Meanwhile, the scattering method calculates OD by measuring the level of light scattered by a sample (in which case the light source is at approximately 90° to the detector), which has been used in other experimental platforms.[8,12] Since Chi.Bio is capable of both approaches we will compare their relative effectiveness in our platform in this note. That said, we use the laser absorption method for OD measurement in the other parts of this manuscript, unless otherwise specified. It is also worth noting that OD measurements only correlate linearly with cell concentration if certain requirements are met (generally best agreement is found at small OD values, and measurements are cell-size dependent), and that this can lead to significant variability between measurements taken by many common platforms used in life sciences research.[13]

##### Laser measurement of OD (Absorption)

For absorption measurements of OD a raw measured value ($R$) is transformed into an initial (logarithmic) OD value ($OD_i$) using the equation:

$$OD_i = log_{10}\Big(\frac{OD_0}{R}\Big) \tag{S1}$$

where $OD_0$ is a calibration value which reflects the measured transmitted intensity through a test tube containing only media. The parameter $OD_0$ conveniently combines scaling factors arising from both the absolute intensity of the light source, as well as its degree of attenuation when passing through the test tube and experimental media. Due to device-to-device variability in absolute laser intensity the value of $OD_0$ needs to be calibrated for each Chi.Bio reactor. Examples of raw measurements ($R$) and $OD_i$ values (calculated using (S1)) are presented in the left column of Fig. S10a,b; the initially differing values align

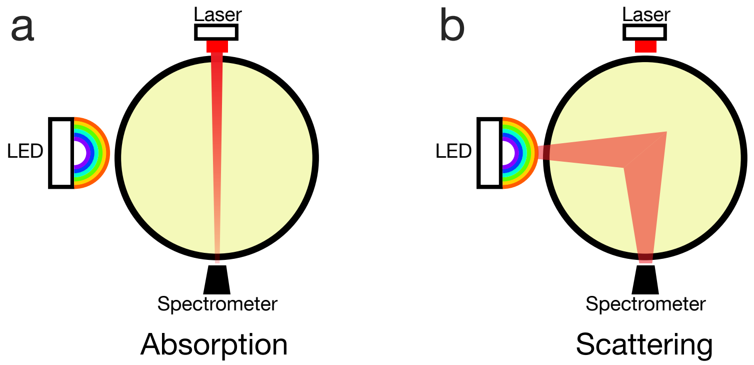

Figure S9: **OD Measurement Theory.** **(a)** To measure OD using absorption a light source is oriented antiparallel to the detector, and light is passed through the sample. **(b)** To measure OD using scattering a side-incident light (in this case at 90° to the detector) is used to illuminate the sample.

closely once $OD_i$ is calculated, compensating for variations in laser intensity.

Values of $OD_i$ can then be mapped to spectrophotometer measurements of OD (in appropriate units of $\mathrm{cm}^{-1}$) by fitting a quadratic calibration function (with no constant term). This functional form is employed to compensate for off-axis scattering events, which become significant at higher optical densities.[13] Such a function is fit in Fig. S10c, which relates the measured values according to:

$$OD\,(\mathrm{cm}^{-1}) = 1.374 \cdot OD_i^2 + 0.3974 \cdot OD_i \tag{S2}$$

When this correction is applied we observe that Chi.Bio- and spectrophotometer-measured OD values align closely, and are linear over two orders of magnitude (Fig. S10d).

**LED measurement of OD (Scattering)**

For scattering measurements of OD a raw value ($R$) measured for scattering from any of the LEDs is transformed into an initial OD value ($OD_i$) using the equation:

$$OD_i = \frac{R}{P_0} - M_0 \tag{S3}$$

where $P_0$ is a LED power compensation factor, and $M_0$ is a media compensation factor. Unlike for absorption measurements of OD (where $OD_0$ was used) these two parameters do not multiplicatively combine; the total raw value must first be divided by the (device specific) raw power level $P_0$ (caused by a combination of variability in LED output and spectrometer sensitivity), and then the (device independent) baseline scattering due to the media/test tube $M_0$ must be subtracted. We measure $P_0$ when no media is in the reactor, and as such it doesn't account for attenuation of the LED's output power as it passes through the media. Fig. S10a,b demonstrate this process for measurements taken using the 623nm LED (middle column) and 6500K LED (right column).

Values of $OD_i$ can then be mapped to spectrophotometer measurements of OD (in appropriate units of $\mathrm{cm}^{-1}$) by fitting a linear calibration function. Such a function is fit in Fig. S10c, which relates the measured values according to:

$$\begin{aligned} OD\,(\mathrm{cm}^{-1}) &= 0.547 \cdot OD_i \qquad (623nm\,LED) \\ OD\,(\mathrm{cm}^{-1}) &= 0.498 \cdot OD_i \qquad (6500K\,LED) \end{aligned} \tag{S4}$$

As for the absorption measurements, when this calibration is applied we observe close agreement between Chi.Bio and spectrophotometer measurements (Fig. S10d).

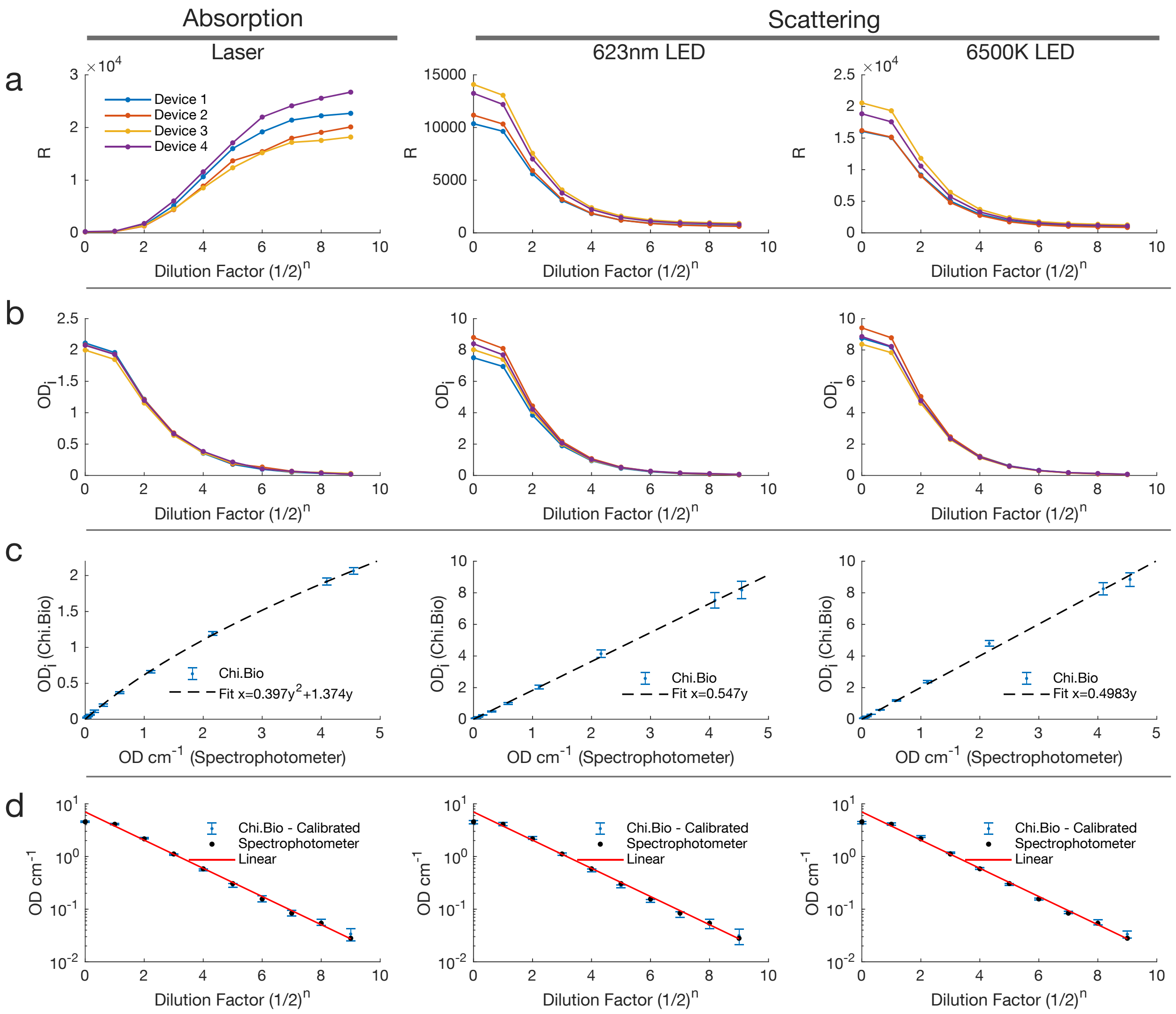

Figure S10: **OD Calibration - Clear Media.** Calibration of absorption and scattering approaches to OD measurement across four devices using standards created by sequential dilutions of evaporated milk in water. This is intended to emulate measurements of cells grown in clear media types. **(a)** Raw readings from the device for each source. All measurements use the Clear spectrometer filter, x1 Gain for the laser and x32 gain for LEDs, and power levels of 0.5 and 0.1 for laser/LEDs respectively. Absorption measurements increase with dilution (as more light can pass through a dilute sample), whilst scattering measurements decrease (less light is scattered in a dilute sample). **(b)** Calculated $OD_i$ values as described in the text. **(c)** Fit of $OD_i$ values to OD ($\text{cm}^{-1}$) values measured by a GeneQuant 1300 laboratory spectrophotometer. **(d)** Plot of calibrated Chi.Bio-measured OD and spectrophotometer-measured OD, demonstrating linearity over two orders of magnitude.

##### Comparing Methods

An obvious question is which of these OD measurement methods in Chi.Bio is more effective. To investigate this we perform a second round of calibration experiments, in this case diluting evaporated milk in LB media (Fig. S11), rather than in water (as in Fig. S10). In both cases new calibration values are required; $OD_0$ values are scaled down due to the additional attenuation by the LB media, and the $M_0$ parameters must similarly be adjusted as the different media have changed scattering properties. In practice $M_0$ does not change significantly between media types (in our case, by $< 5\%$ when moving between water and LB), though this small difference has a significant impact when cells are at low densities. Our results demonstrate that the previously fit calibration functions (from Fig. S10c) match more closely for the Laser-based absorption measurements when compared to scattering measurements (Fig. S11c). We hypothesise that this error in the LED measurement is due to absorption of the LED light as it passes through the LB media, which is captured by the coefficient in (S4). This factor is better compensated for absorption-based measurements, since it is multiplicatively included in the value of $OD_0$. Therefore, if precise OD readings are required the absorption approach to OD measurement in Chi.Bio may generally be favourable if different media are used, though better yet is re-calibration with each new media type employed. That said, even without re-calibration the measurements made by both techniques remained close to the spectrophotometer readings when LB media was substituted for clear media (Fig. S11d). An additional consideration is that absorption measurements can utilise light sources at far lower optical power, thereby minimising their impact on light-sensitive biological systems (i.e. optogenetic circuits). This is observed in Fig. S10, where a 32x increase in gain is required for scattering measurements to achieve a similar absolute magnitude (when compared to absorption measurements), despite the LED's optical power output being at least an order of magnitude greater than the laser's at the set-points chosen.

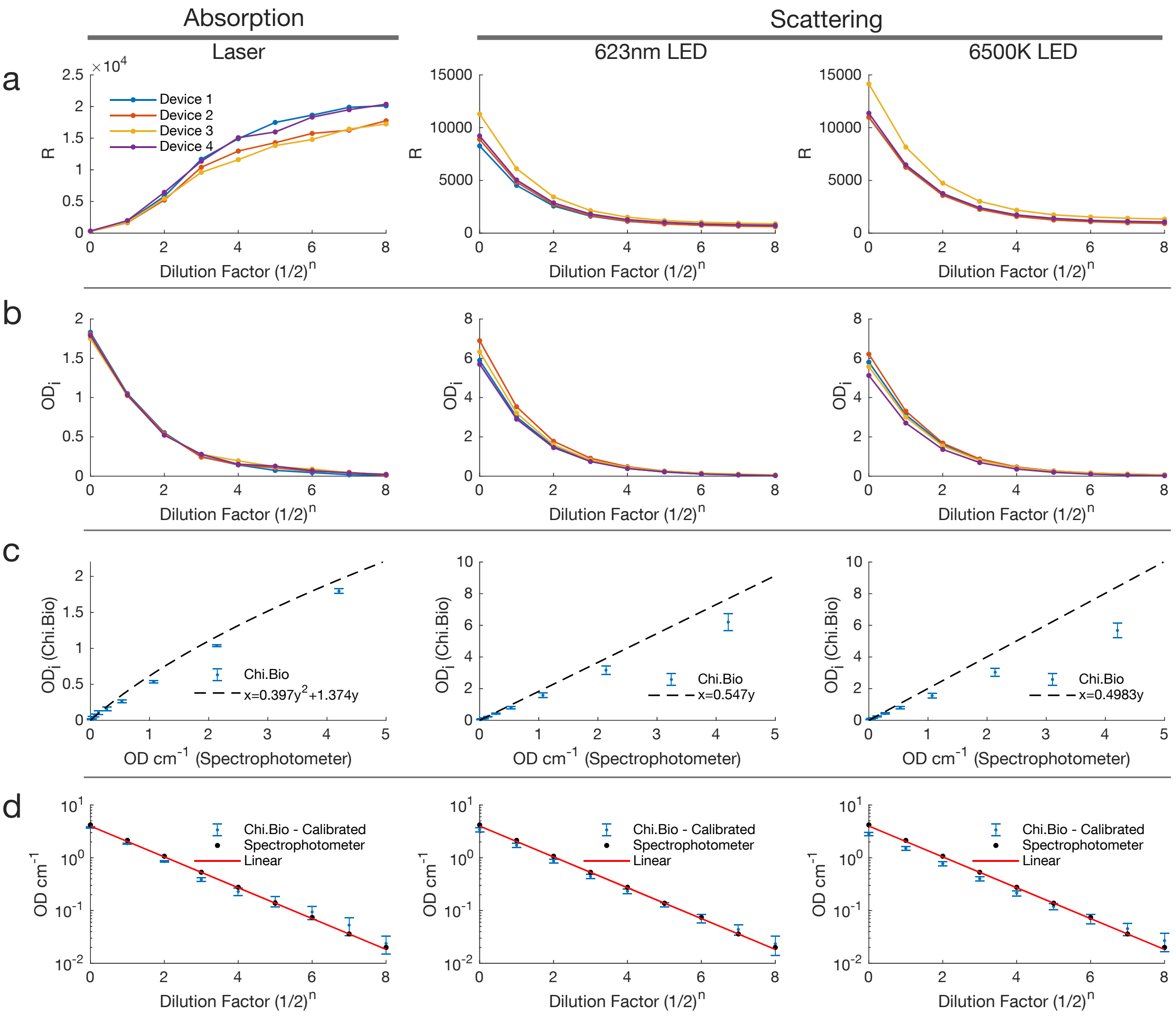

Figure S11: **OD Calibration - LB Media.** Calibration of absorption and scattering approaches to OD measurement across four devices using standards created by sequential dilutions of evaporated milk in LB media. Device 4 differs to that used in Fig. S10. **(a)** Raw readings from the device for each source. All measurements use the Clear spectrometer filter, x1 Gain for the laser and x32 gain for LEDs, and power levels of 0.5 and 0.1 for laser/LEDs respectively. **(b)** Calculated $OD_i$ values as described in the text. **(c)** $OD_i$ compared to previous calibration function fit in Fig. S10c. **(d)** Plot of calibrated Chi.Bio-measured OD (using calibration function fit in Fig. S10c) and spectrophotometer-measured OD.

### Note S6 Laser Analysis

The 650nm OD laser (and its driving circuitry described in Note S3) is designed to provide stable measurements of optical density as conditions vary. Once calibrated (Note S5) ideally OD readings should have minimal noise, and be invariant as the temperature in the reactor changes.

In Fig. S12a the measured output power of the laser is plot against the power set-point for four devices. Some variability is present between diodes, which we hypothesise arises due to differences in the relative geometry of the laser source and its internal photodiode. All devices turn on at an input set-point of $\approx 0.1$ V due to the 2 MΩ resistor in the driving circuit, which is used to compensate for the OpAmp's bias voltage (without this resistor the laser may not be completely off when $V_{in} = 0$). In general the shape of these curves is not of great significance since we do not vary the laser intensity during operation, and calibrate (as in Note S5) with respect to the absolute output at a constant power set-point. Fig. S12b presents the measured spectrum of the laser's output across four diodes, which suggests that the laser's output wavelength may be (according to our spectrometer) slightly lower than expected (as the 620nm measurement is much larger than that at 670nm, which is closer to the laser's specified wavelength peak of 650nm).

Fig. S12c analyses noise in the laser's OD measurement of a constant target. We observe that there is minimal inter-shot variability arising in the laser/spectrometer themselves, with the vast majority of noise in the OD measurement arising with the onset of stirring. Note that in this case stirring is active for 40 seconds out of each minute, with 10 seconds given for the liquid to settle prior to measurement (as in automated experiments as described in Fig. 2g). Nevertheless, active agitation of the liquid results in moving of spatial inhomogeneities in culture density which are picked up by the sensor. If very precise readings of OD are required the impact of culture inhomogeneity can be minimized by averaging each measurement over a longer time period.

Secondary photodiodes can be used to mitigate the impact of temperature fluctuations on OD readings.[9,10] Our system implements similar functionality directly in its hardware, as illustrated in Fig. S6a. Fig. S12d analyses the stability of this system by measuring a stable solution's OD as it is heated from 25 $°C$ to 47 $°C$. We observe a small degree of variability with respect to temperature (which we hypothesise may be caused by varying photodiode alignment/output with temperature), though this is of an order comparable with the anticipated stirring-induced noise in OD measurements.

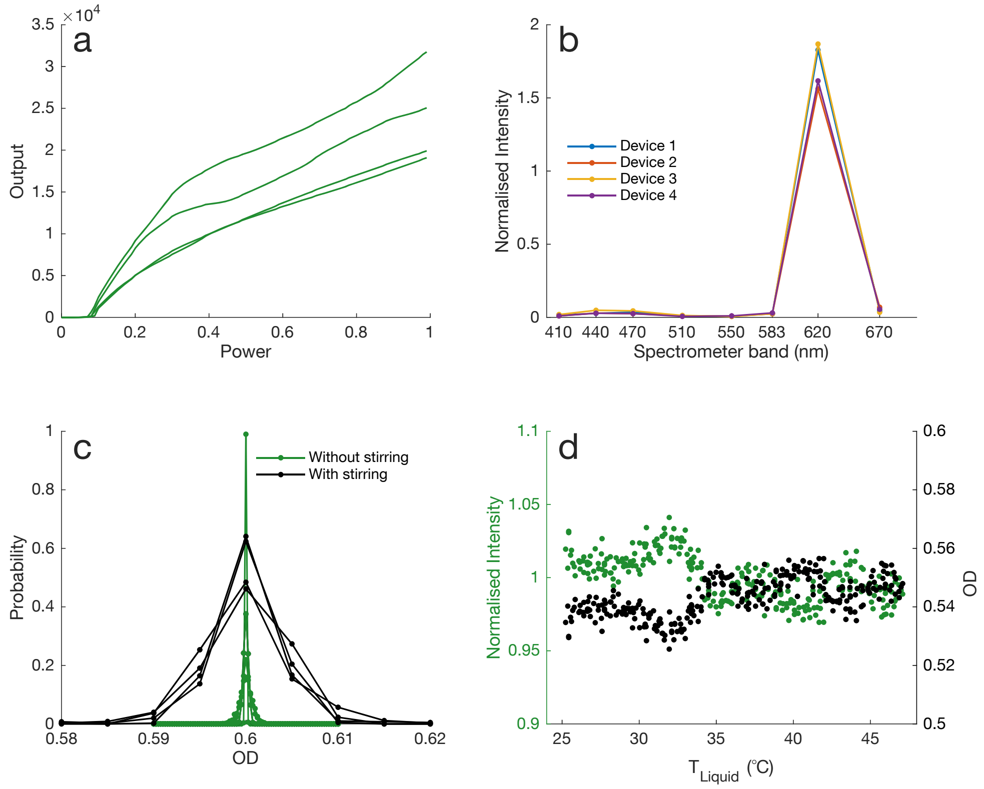

Figure S12: **Laser Analysis.** **(a)** Measured laser intensity (with Clear spectrometer filter) vs power set-point for four devices. **(b)** Spectral decomposition (normalised by Clear filter measurement) for lasers in four devices. **(c)** Probability density function of measured OD of evaporated milk standard solutions for four devices, measured each minute for six hours. All standards had OD between 0.55 and 0.65, and have been re-scaled here (to a mean of 0.6) to directly compare distributions. For the stirring experiment the solution was stirred for 40 seconds of each minute, and then allowed to settle for 10 seconds prior to measurement. **(d)** Measured laser intensity normalised by mean (left axis) and calculated OD (right axis) for a MacFarland standard solution of varying temperature. The temperature of the solution was linearly increased over the course of 5 hours from 25 $°C$ to 47 $°C$, with measurements taken each minute (and proceeded by 40 seconds of stirring to prevent settling).

### Note S7 LED/Spectral Analysis

The visible LED and spectrometer are designed to provide stable measurements with minimal variability between conditions or devices. In this note we examine the performance (and in particular, variability) of these components as the output power level, temperature, or device changes.

Fig. S13 plots the measured intensity across the spectrum for each visible LED as its power set-point is changed. Ideally this relationship would be linear, thereby enabling experimental applications that require predictably tunable light intensities (e.g. optogenetics). In such cases the absolute LED output levels can easily be normalised between devices (i.e. as is done in Fig. S13) to minimise variation in absolute light intensity between reactors. We observe that all LEDs are almost perfectly linear up to a power level of $\approx 0.5$ (which corresponds to an optical output significantly higher than needed for almost any application), and that some (i.e. the 595nm LED) are more sensitive than others above this point. In general the spectral output of each LED has minimal variability with respect to its set power level, which is illustrated in Fig. S14. The component which exhibits the most change is the 500nm LED, which shifts slightly toward a higher wavelength peak as its output power increases (Fig. S14c).

The spectral breakdown of each LED at a fixed power level is plot for four devices in Fig. S15. Each measurement is normalised by the spectrometer's Clear filter intensity (as done for fluorescent protein measurements, described in Note S15); when this is done we observe minimal inter-device variability in the LED spectrum. This is the metric of significance for noise in fluorescent protein measurements made by Chi.Bio: The minimal variability in each measurement band translates to minimal inter-device variability in fluorescence measurements.

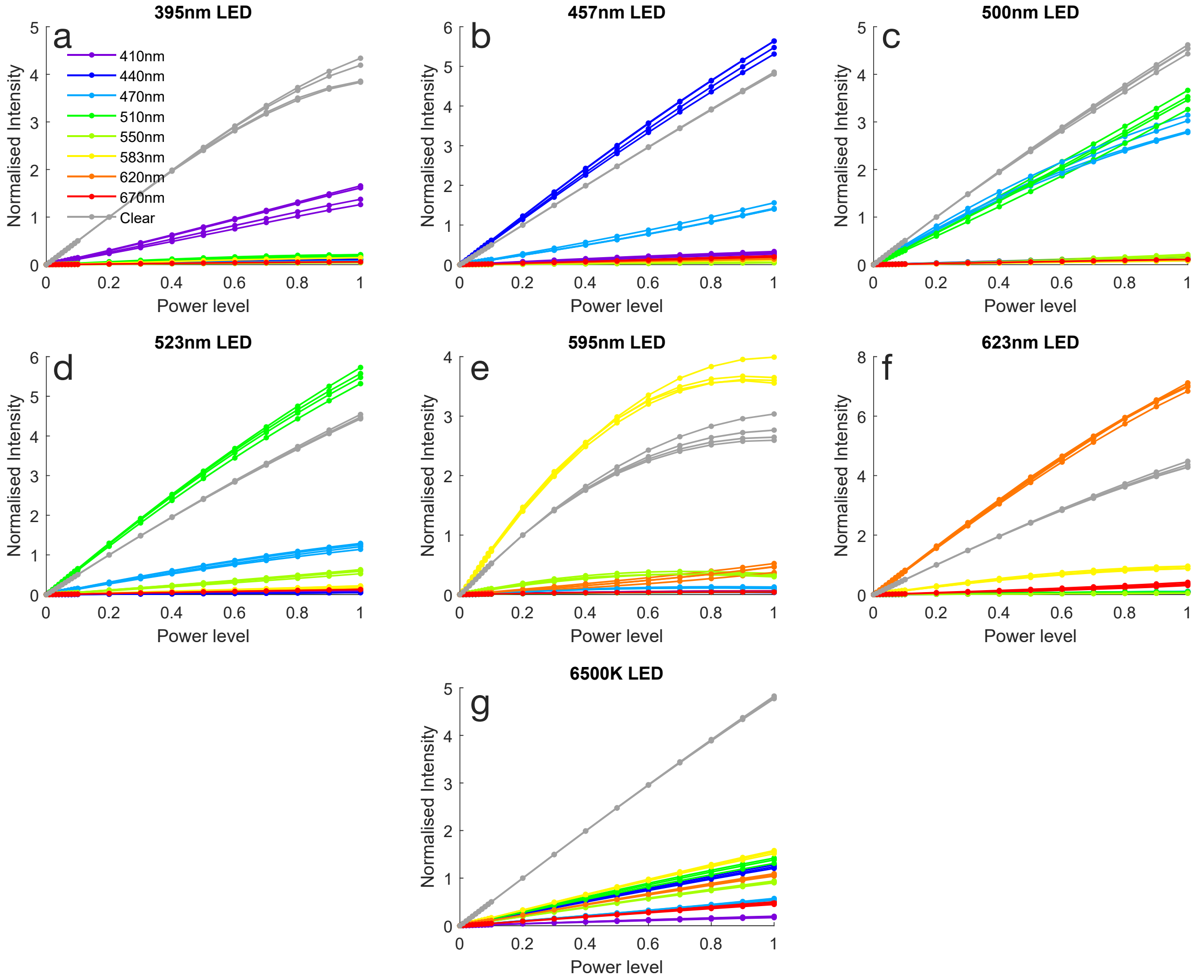

Figure S13: **LED Output intensity vs Power Set-point.** Each LED's measurements are normalised by the intensity measured by the Clear spectrometer band when power level is 0.2. Data presented is for four devices. Examples of an LED at different power levels is presented in Fig. S7.

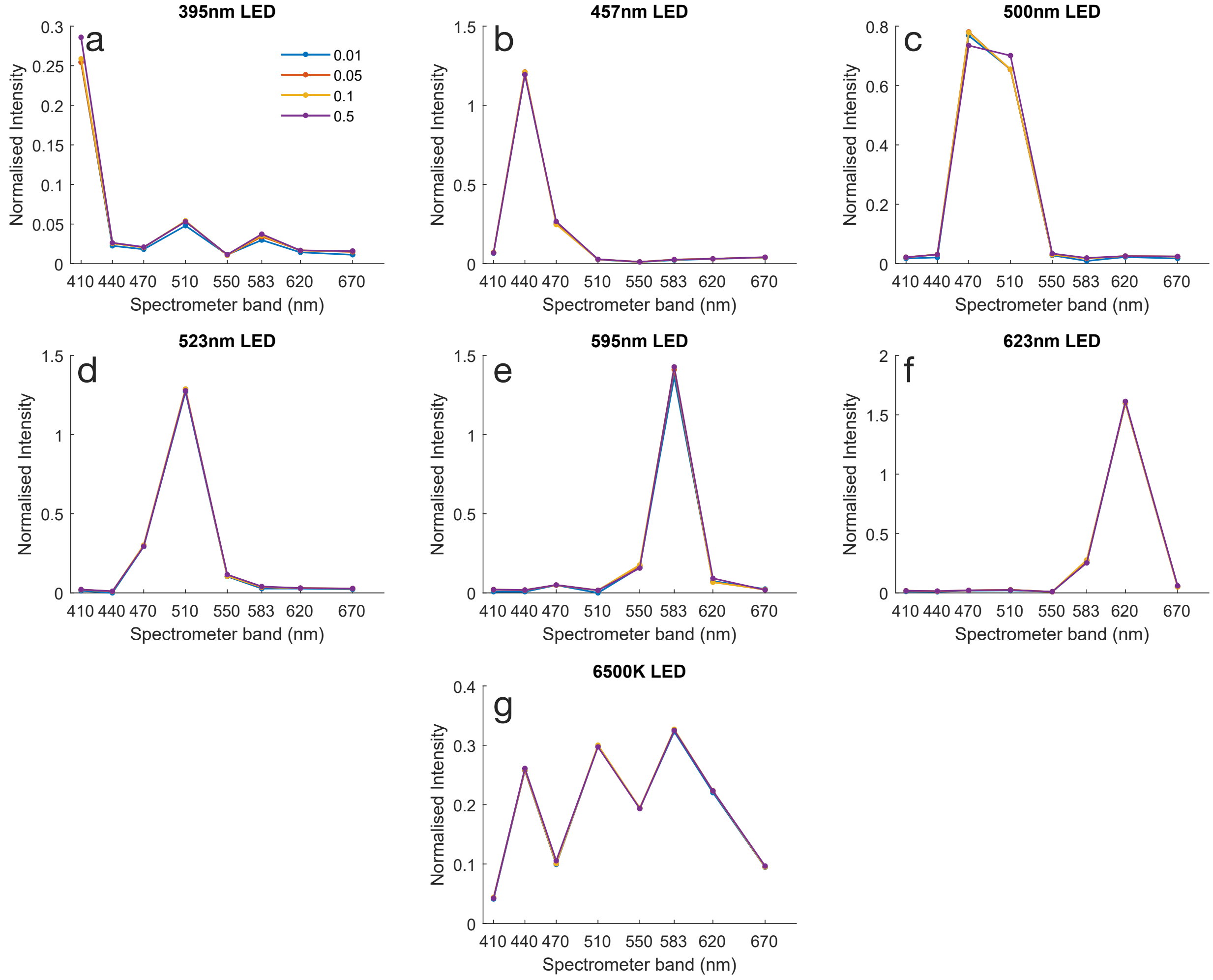

Figure S14: **LED Spectrum vs Power.** Spectrum of each LED output as its power set-point varies, normalised by the intensity measured by the Clear spectrometer filter.

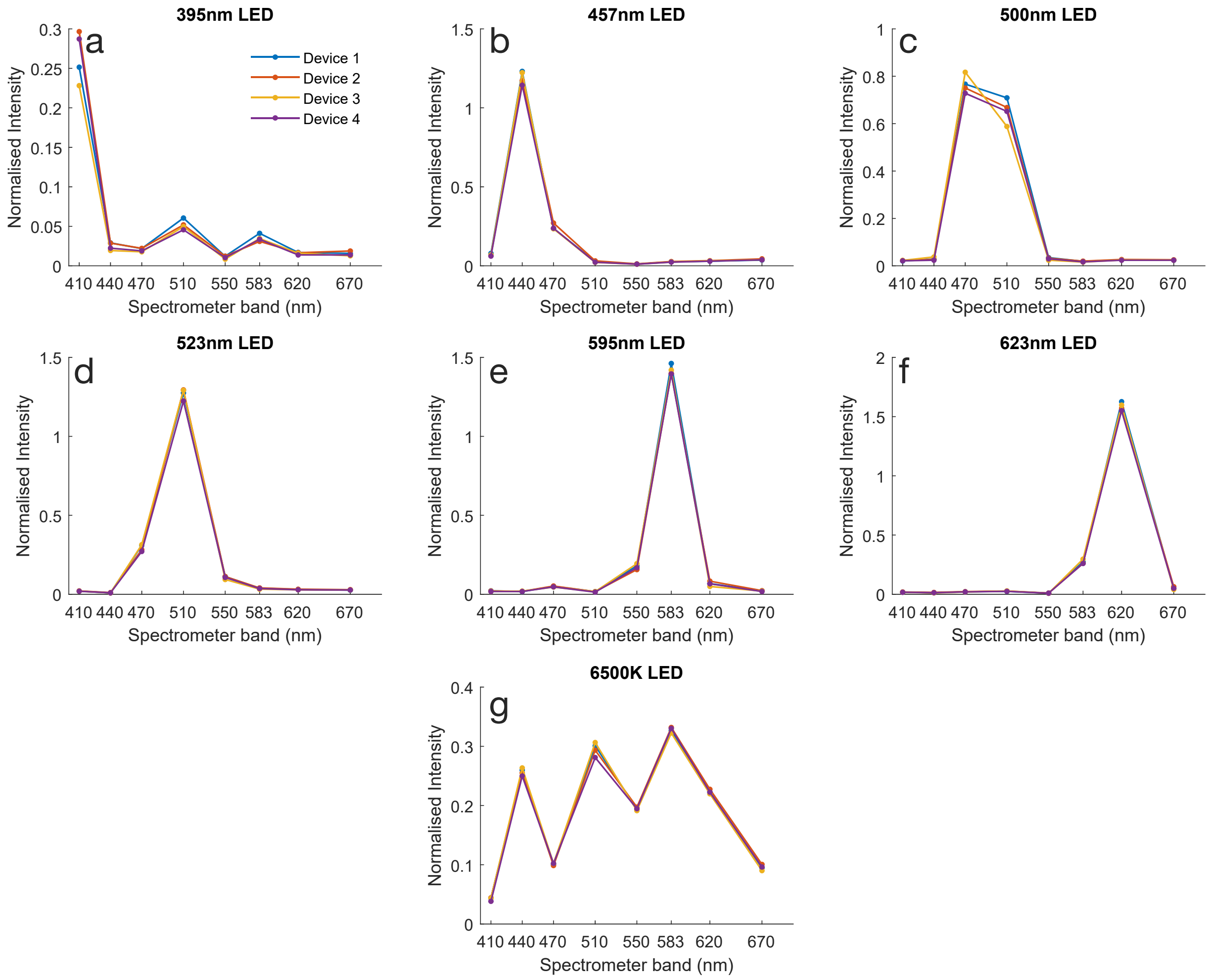

Figure S15: **LED Spectrum vs Device.** Differences in spectrum of each LED output (at power level 0.1) for four different devices. The measurements in each wavelength band are normalised by the intensity measured by the Clear spectrometer filter.

Fig. S16 investigates the dependence of the visible LED's output on temperature as a standard solution is heated from 25 $°C$ to 47 $°C$ (as was done for the laser in Fig. S12). We observe minimal difference in the spectral breakdown or absolute intensity of any LED as temperature changes in this range, with the exception of the 595nm LED for which the output intensity decreases by approximately 0.5% per degree of temperature increase. That said, this level of variability is likely to be insignificant for most biological applications (where media temperature is not varied so greatly), and can be mitigated by adaptively adjusting its set-point to maintain its measured output. The 595nm LED was also observed to decrease in intensity at high power outputs (Fig. S13e), which was likely caused by heating of the diode at high power.

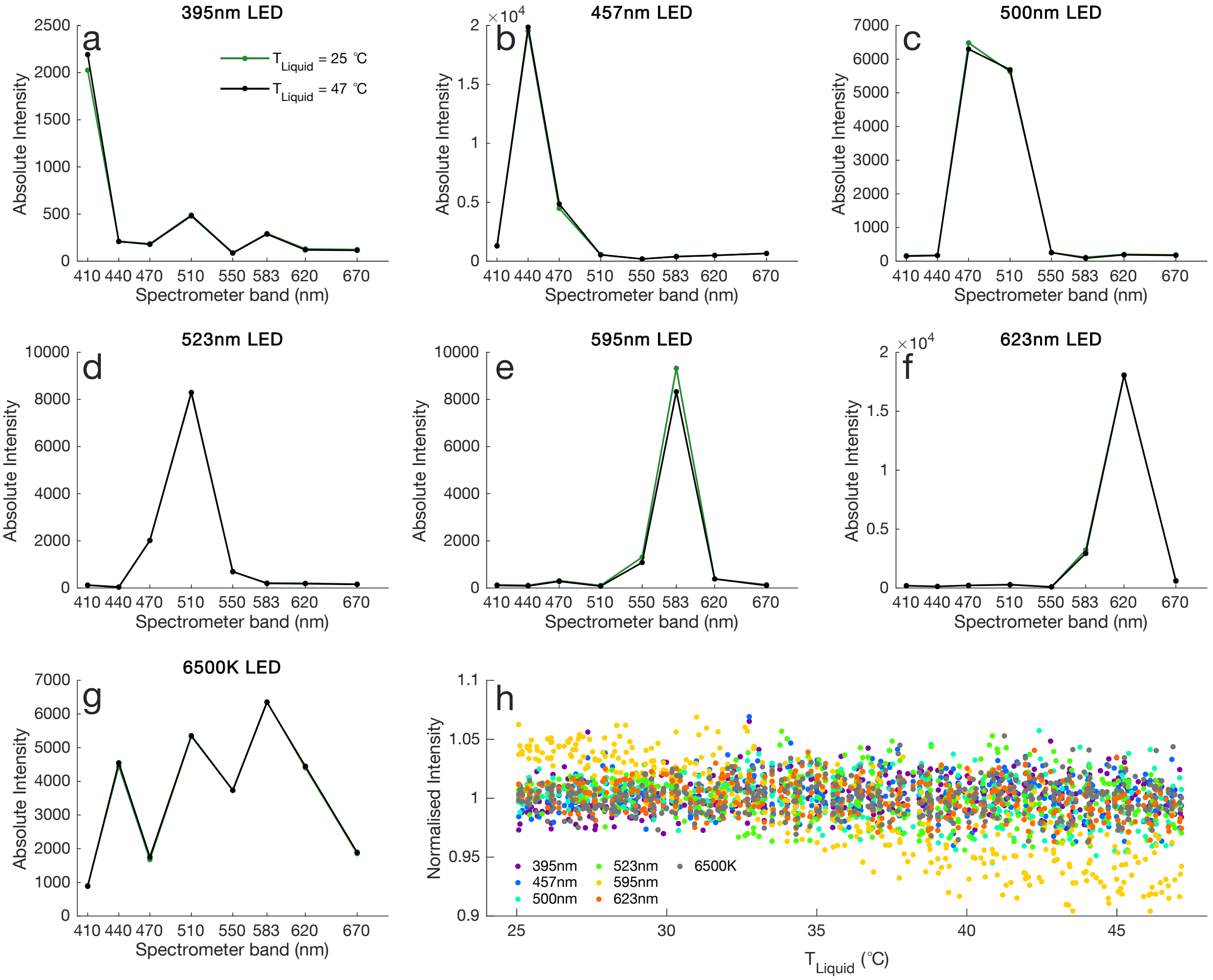

Figure S16: **LED Spectrum vs Temperature.** **(a-g)** Absolute measured spectrum of each LED as a MacFarland standard is heated from 25 $°C$ to 47 $°C$ over five hours (with stirring). **(h)** Output intensity of each LED measured by the Clear spectrometer filter once per minute (and normalised by mean value over the entire experiment).

To analyse the sources of inter-device variability, in Fig. S17 and S18 we take spectral readings of each LED with two devices (Device 1 initially has LED A and Spectrometer A, Device 2 has LED B and Spectrometer B), and then disassemble these devices, switch out their spectrometer chips, and perform the same measurements. We observe that some variability arises following a change in spectrometer (rather than LED): Measurements of different LEDs with the same spectrometer are more similar than measurements of the same LED with different spectrometers. When we normalise these readings by the intensity measured by the Clear spectrometer band (Fig. S18) this variability reduces significantly. This implies that inter-device variability in measured LED intensity is largely determined by the total light reaching the spectrometer (which is varying by a similar factor across all spectrometer wavelength bands). We hypothesise that this variability is caused by a combination of small differences in component alignment (the spectrometer and LED are attached to different PCBs which are bolted together) and variability in spectrometer aperture size (which determines the total light incident upon its sensors). This conclusion contrasts the alternative possibility, that the spectrometer measurement bands differ significantly in their wavelength-filtering profile from device to device, in which case normalisation by Clear filter intensity (i.e. moving from Fig. S17 to S18) would have had minimal impact on variability.

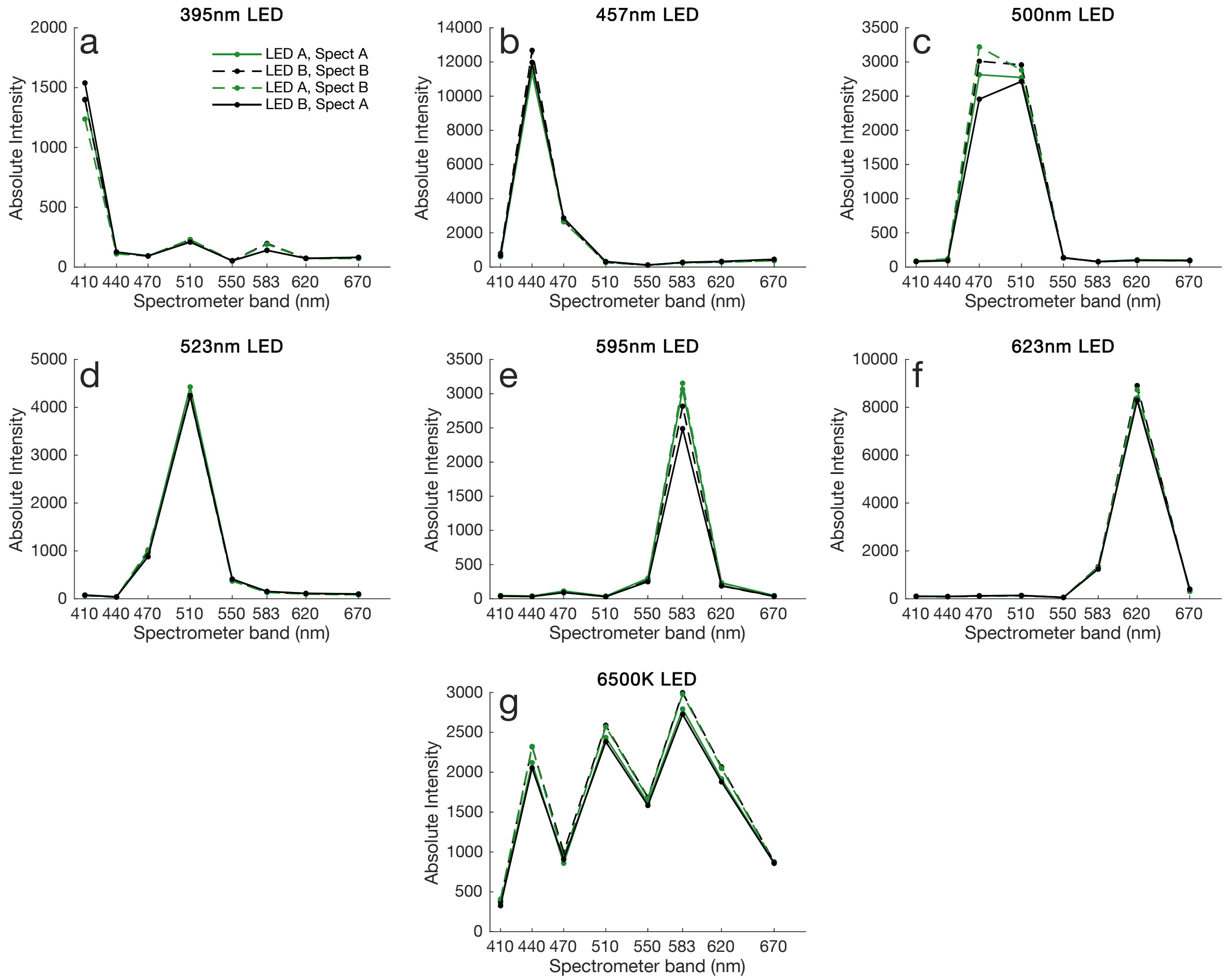

Figure S17: **Absolute Inter-Component Variability.** Absolute measured spectrum of each LED for two Chi.Bio devices before and after switching their spectrometer chips. Device 1 initially has LED A and Spectrometer A, and Device 2 has LED B and Spectrometer B.

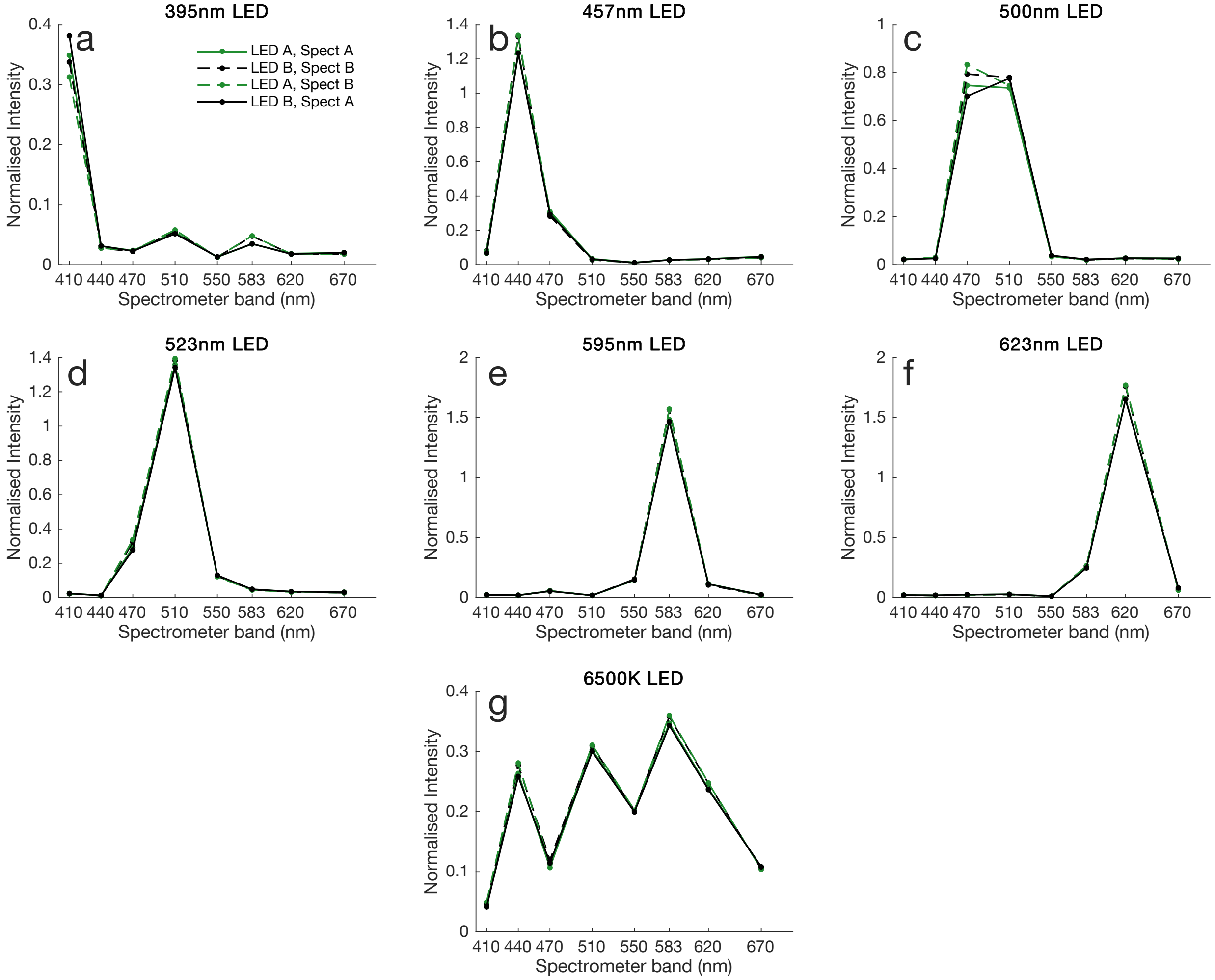

Figure S18: **Normalised Inter-Component Variability.** Similar to Fig. S17, but with all spectra normalised by the LED intensity measured by the Clear spectrometer filter.

### Software and Automation

#### Note S8 Operating System

Chi.Bio's operating system comprises a HTML/Javascript front-end for user interactions, and a Python back-end which provides the underlying processing and control functions of the platform. In this note we briefly summarise the structure and function of each of these components. All software is open source and made available on our project GitHub (https://github.com/HarrisonSteel/ChiBio) to allow ongoing community contributions.

##### Front End - HTML/Javascript

The front end of the Chi.Bio operating system is responsible for direct interactions between users and the platform, which happen through an easy-to-use web interface (discussed in Note S9). The structure of the UI web page is implemented in HTML, which defines the locations/sizes/text of each control or input field. If a control input is changed, or new data become available, each entry in the HTML page is updated using a Javascript program which assigns values to each HTML element according to data output by the Python back-end. The Javascript component is also responsible for defining the state (active/inactive) of the various buttons on the interface, as well as disabling (greying-out) those which are not usable at a given time. Data is prepared for plotting by the Javascript component, taking advantage of the free-to-use Google Charts toolbox. When a control in the HTML page is activated a command (and accompanying input data if appropriate) including this information is sent (through the web server) to the Python back end.

##### Back End - Python

The processing, data-handling, and control features of Chi.Bio are all implemented in a Python back-end script which operates over a number of levels of abstraction.

*Digital Level:* At the lowest level of this system is the exchange of digital communications with each of Chi.Bio's sub-systems via the two-wire $I^2C$ bus. As described in Note S2, these signals are multiplexed to communicate with up to eight connected reactors. Multiplexing is implemented in the operating system using a custom bus-handling protocol which (when a $I^2C$ command is requested) takes command of a digital lock on the $I^2C$ bus, switches the multiplexer to connect the Beaglebone's $I^2C$ lines to the appropriate reactor, executes the read/write $I^2C$ command required, and then disconnects the multiplexer. During this process the outcome of each data transfer is checked, and errors (e.g. if a bit is lost along the way) are managed. Using this bus-handling system $I^2C$ read/write commands can run at up to $\approx$500Hz, which provides ample bandwidth for eight devices connected to the single Beaglebone $I^2C$ controller.

*Library Level:* At the next level are a range of library functions which govern the digital interfacing with each sub-system. These include (for example) commands for spectrometer setup and data-read, or PWM driver setup. Such commands make it very straight-forward to implement new features in the operating system, since users can specify (for example) which spectrometer bands they would like to measure (and with what gain), and the library functions automatically execute the many digital $I^2C$ commands required to achieve this.

*Automation Level:* Automation functions (e.g. for pump modulation, OD/temperature regulation, or the overall experimental cycle) run in Python threads, allowing may processes to occur concurrently during the platform's operation. This is essential to the parallelisation of data processing and handling which is required to run multiple reactors (each with different experiments) from the same control computer. Each thread carries the index of the reactor with which it is interfacing, though it is also possible to access data from / send data to other reactors running in parallel.

*Data Level:* As data is input/measured it is stored in data arrays (unique to each connected reactor), which maintain a record of all control parameters, measured values, and actuation inputs that arise during an experiment. In each automated experimental cycle measured data from that cycle is appended to an external CSV file for permanent storage, and the platform's setup information is recorded (in JSON format) in a text file (which can easily be read as a complete data structure into other programming languages).

*Input/Output Level:* Data is exchanged between the user and the Python interface via a web server implemented using Flask[14] and Gunicorn.[15] This web server packages data and setup parameters into JSON structures, which are then processed by the Javascript component of the Front End (described above). The web server's interface (i.e. the HTML page) can be accessed by navigating (in a web browser) to its IP address on a connected PC or network.

### Note S9 User Interface

The top of the UI (Fig. S19) includes many controls for running and customising experiments on Chi.Bio. In the top left is a device switchboard; clicking on a different device switches the interface to display its setup parameters/data. The rest of the interface includes controls to (for example) calibrate OD measurements, set the turbidostat OD target, adjust LED powers, manually control pump speeds and direction, or adjust the temperature set-point. When a particular item is in use/enabled its switch goes blue to indicate this, and buttons are greyed out in circumstances when the platform's automation functions are using that sub-systems (e.g. the switches used to set new rates for Pump 1 and 2 are greyed out in Fig. S19 because the OD regulation algorithm is controlling them). Most hardware outputs (e.g. optical outputs, stirring) can have their power/speed set in a range from $[0, 1]$, where 1 indicates maximum power and 0 is off. The pump outputs can be set in the range $[-1, 1]$, where negative values indicate reversed direction. Though light/thermometry measurements are usually taken automatically during an experiment, the UI includes "Measure" buttons for each of these systems if the user wants to take readings outside of the automated modes.

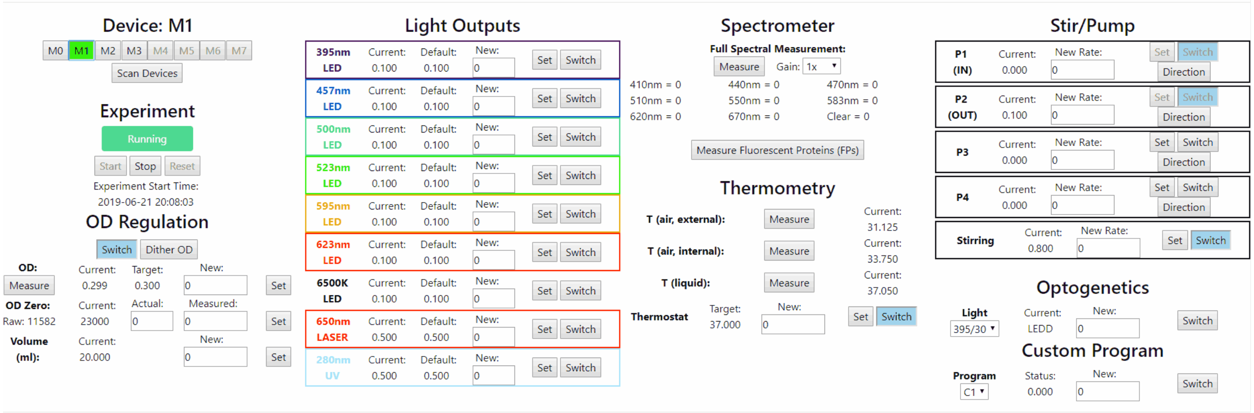

Figure S19: **UI Controls.** The top of the web user-interface, which can be accessed from the user's web browser. For a detailed explanation of each button see the operation manual provided on the Chi.Bio website (https://chi.bio/operation/).

The lower part of the UI (Fig. S20) is primarily used for plotting data as they are collected. Different figures present the current/target OD, current/target temperatures, pumping rates, and each fluorescent protein measurement that is in progress. These figures are updated each minute as new data arrives. The data that is displayed in these figures is interpolated to lower resolution each $\sim 3$ hours in order to minimise the size of files being handled by the user's web browser. Fluorescent protein measurements are also set-up and enabled in the lower component of the UI. Five options are provided for each measurement (as well as an "Activate" switch to enable it): the excitation wavelength, the baseband (see Note S15), first and second emission wavelength bands, and the measurement gain.

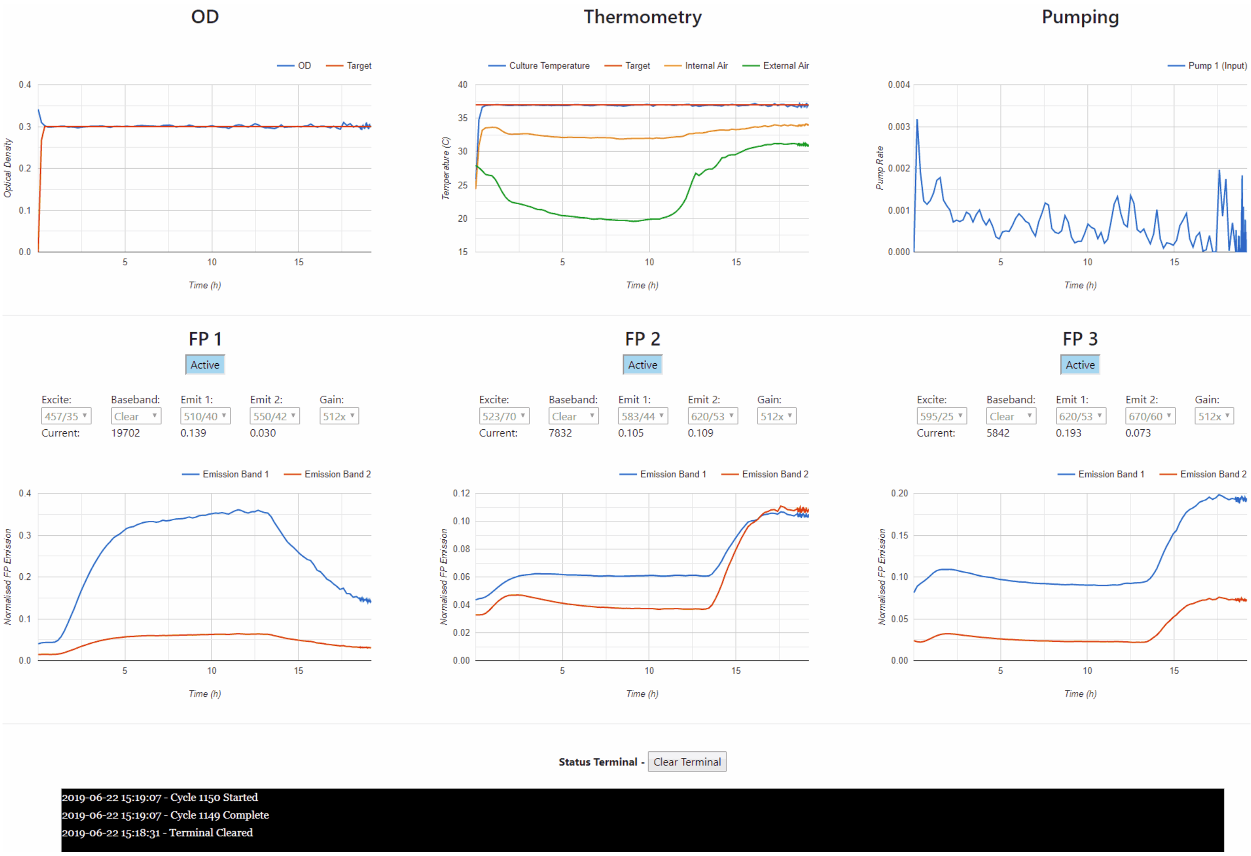

Figure S20: **UI Data Plotting.** The bottom of the web user-interface, which presents data in real-time as it is collected.

### Note S10 Customisation

In practice many more experimental procedures than those described in this paper can be performed automatically on Chi.Bio. These can be implemented in the Python back-end of the operating system, for which we have developed a basic “custom program” framework (with controls in the bottom-right of Fig. S19).

Inside a custom program users can write their own functions that combine measurements from one (or many) devices with user-inputs or outputs from other processes running on the control computer. These can be used to control any combination of the platform’s actuation outputs using the built-in library functions and any available Python library installed on the control computer. An example of a custom program developed in this study is the optogenetic feedback controller described in the main text, which uses measured fluorescence values to calculate new optogenetic LED intensities, and then switches these LEDs on for part of the experimental cycle.

For a detailed guide to implementing new software functionalities see the Chi.Bio GitHub (https://github.com/HarrisonSteel/ChiBio).

### Note S11 Temperature Regulation

A complex feedback control algorithm uses temperature measurements (taken by the infrared thermometer) to regulate the heat input (provided by the heat plate) to control the temperature in the growth chamber. The control algorithm employed also takes into account the inflow rate and temperature of fresh media being pumped into the device (assumed to be at room temperature, measured by the thermometer on the control computer). The control algorithm comprise two main components:

**Proportional - Integral (PI) Controller:** The PI controller uses the current temperature error, $e = T - T_t$ (where $T$ is the measured temperature and $T_t$ is the desired target temperature) to drive the media temperature toward its intended target. This controller operates in two modes; when $|e| > 2.0$ a large proportional gain is used for fast convergence, and when $|e| < 2.0$ a smaller proportional term as well as an integral term is employed to minimise steady-state error.

**Model-Predictive Controller (MPC):** In addition to the PI controller a simple model-predictive controller is included, which applies additional heat at a rate proportional to $P_1(T_t - T_M)$ (where $P_1$ is the rate of media input, and $T_M$ is the input media temperature). This model-predictive component is used to minimise temperature undershoot when cold media is pumped into the device for a short period of time.

The combined controller enables temperature regulation which is precise (the culture remains close to its set-point), fast-responding (a new temperature set-point is rapidly converged upon), avoids overshoot/undershoot of the set-point, and is robust to environmental fluctuations (e.g. addition of liquids of a different temperature to the growth chamber). This is demonstrated in Fig. 2h of the main text, and Fig. S21 below (where the accuracy of the control algorithm is analysed).

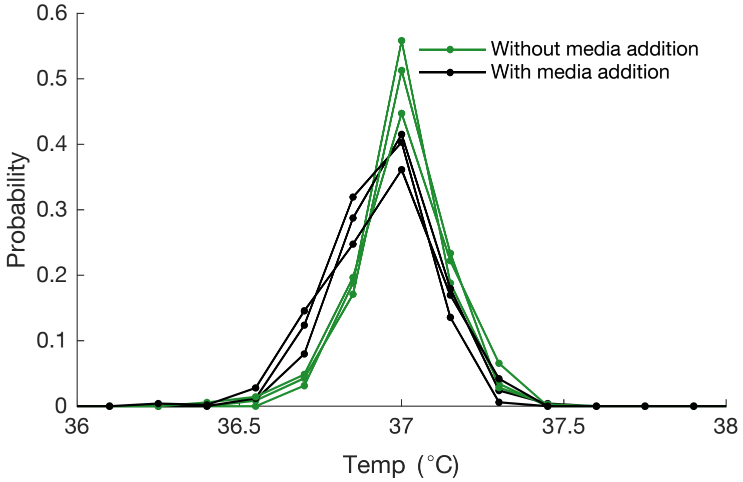

Figure S21: **Growth Chamber Temperature Distributions.** Probability distribution of temperature in the growth chamber when the thermostat is set to 37 °C over the course of 8 hours (for three different devices). In both cases ambient temperature is ≈ 26 °C. For the "With media addition" experiment *E. coli* is being grown in turbidostat mode, necessitating continual input of fresh media (which is at room temperature). Consequently, the temperature distribution is slightly skewed toward lower temperatures as the input of (cold) fresh media temporarily reduces the growth chamber's average temperature. The MPC controller (described above) largely mitigates this effect.

### Note S12 OD Regulation

Cell density/OD regulation (turbidostat functionality) is implemented by a control algorithm which measures the culture's optical density using the laser (or LED, see Note S5) and spectrometer, and actuates OD change (via dilution with fresh media) using the pumping system. To determine the appropriate rate of fresh media input we employ a PII (Proportional-Integral-Integral) controller which operates on the error between the current ($OD$) and target ($OD_t$). The PII controller terms have gains of $G_p$, $G_{I1}$, and $G_{I2}$ respectively, where $G_{I2} \gg G_{I1}$. The P and first I term are primarily responsible for OD regulation near to $OD_t$. The second integrator is set to zero any time that $OD < OD_t$; its purpose is to account for rare situations that may arise if the tubing of the input peristaltic pump is not set up correctly, causing it to have a temporarily reduced suction and subsequent reduction in the rate of media addition. The output pump rate (i.e. removal of waste media) is set significantly higher than the input pump rate at all times to guarantee a much greater rate of media removal, ensuring the volume of media in the growth chamber does not rise. Thus most of the time the output pump will be pumping air out of the growth chamber. Fig. S22 illustrates the accuracy of the OD control algorithm during exponential growth of *E. coli*.

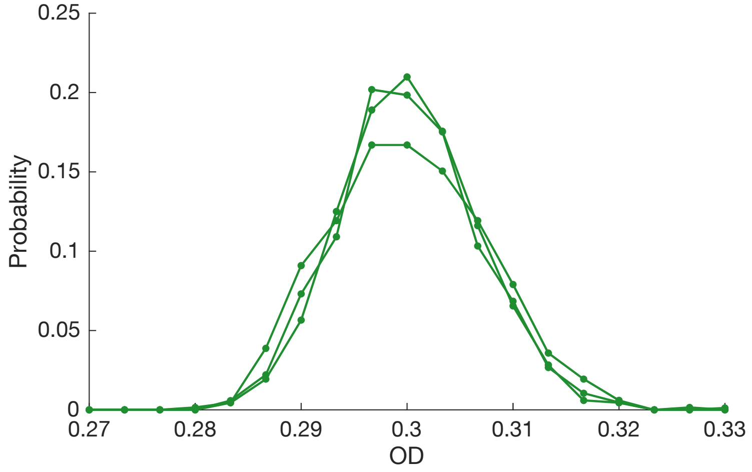

Figure S22: **OD Distributions.** Probability distribution of measured OD (using the OD Laser) when the control algorithm is set to maintain *E. coli* at an OD of 0.3 over the course of 10 hours (for three different devices).

### Experimental Notes

#### Note S13 UV Recovery

In Fig. 3c we found that a cell colony was not entirely killed by intermediate UV levels. Fig. S23 demonstrates that a colony can adapt to UV dosing, with growth rate recovering to the same level achieved in the absence of UV.

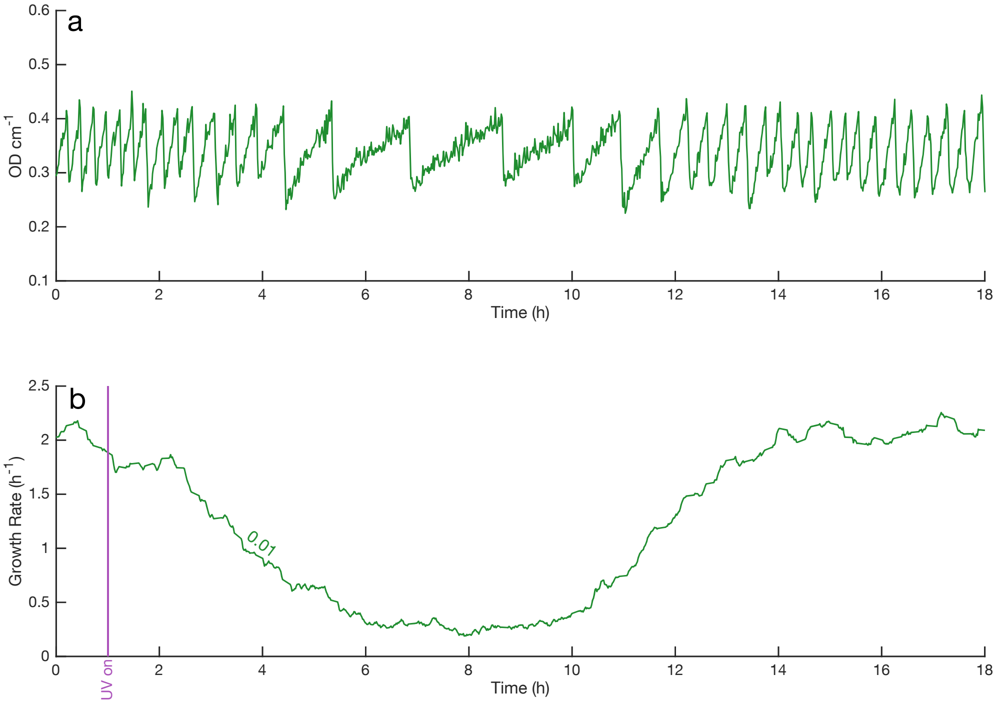

Figure S23: **Response of growth to intermediate UV dosing.** **(a)** OD profile during the experiment, set to follow a dithered pattern for estimation of growth rate near a constant cell density. Growth initially slows when UV is enabled (which occurs after 1 hour, at intensity 0.01), but recovers over time. **(b)** Growth rate calculated from OD data in panel **a**, demonstrating that the total growth rate is reduced by $\approx 85\%$ before returning to a similar growth rate to that measured prior to UV activation.

### Note S14 Biofilm Measurements

The study of biofilms has broad implications for medical research and industrial biosynthesis, with *E. coli* frequently used as a model organism for analysis of biofilm formation on abiotic surfaces.[16] A number of experimental methods exist for measurement of biofilm formation and growth (spanning batch- and continuous-culture implementations), which demonstrate that biofilm accumulation depends on many factors including nutrient availability, strain choice, methodology, surface properties, and shear conditions.[16,17] Chi.Bio's hardware can thus be used in a number of configurations (Turbidostat, Chemostat) to study biofilm accumulation, and more advanced techniques may additionally utilise its spectral measurement capabilities. Here we focus on a simple proof-of-concept turbidostat-based measure of biofilm accumulation, which takes advantage of past work on the modelling of growth in continuous culture regimes.[18]

##### Measurement Theory

We start by analysing the anticipated impact of Biofilm presence on measurements of OD made by the laser (i.e. the absorption method). We express the total light intensity incident upon the spectrometer ($I$) using the Beer-Lambert law:

$$I = I_0 \cdot \exp(-A_c\rho_c) \cdot \exp(-A_b\rho_b) \tag{S5}$$

where $I_0$ is the initial (un-attenuated) intensity, $A_c$ and $A_b$ are absorbance coefficients of cells in media and biofilm respectively, and $\rho_c$ and $\rho_b$ are the densities of the cells in media and biofilm respectively. Note that these quantities will have different units; the biofilm parameters are expressions in terms of area, whilst the cells-in-media parameters are in terms of volume, but this is not critical to the current analysis.

If $I$ in (S5) is to be kept constant (i.e. the platform is in Turbidostat mode) then so must be the quantity $C_0$ in:

$$C_0 = A_c\rho_c + A_b\rho_b \tag{S6}$$

We can observe from this relation that if $\rho_b$ increases (i.e. a biofilm grows on the inside of the test tube) then a smaller density of cells is required to maintain a constant transmitted intensity ($I$, the apparent optical density measured by Chi.Bio). We can take a time derivative of (S6) to yield:

$$\dot{\rho}_c = -\frac{A_b}{A_c}\dot{\rho}_b \tag{S7}$$

where the dot operator indicates time derivative. This can then be combined with a growth equation:

$$\dot{\rho}_c = (\mu_c - P_{in})\rho_c + \mu_s\rho_b \tag{S8}$$

where $\mu_c$ is the specific growth rate of the cells, $P_{in}$ is the input (dilution) pump rate, and $\mu_s$ is the rate of cell shedding from the biofilm (cells which grow on the biofilm and escape into solution[19]). We can combine (S6), (S7), and (S8) to give:

$$P_{in}(t) = \mu_c + \frac{\dot{\rho}_b + \frac{A_c}{A_b}\mu_s\rho_b}{\frac{C_0}{A_b} - \rho_b} \tag{S9}$$

We observe from (S9) that as $\rho_b \to \frac{C_0}{A_b}$ (and correspondingly $\rho_c \to 0$) the required pump rate $P_{in} \to \infty$. Conceptually this situation arises when the biofilm's optical density approaches the Turbidostat set point, requiring an ever-decreasing (and eventually negative) cell density and large corresponding pump rate. That said, the pump rate may also saturate as the numerator in (S9) grows (whilst $\rho_b$ is significantly less than $\frac{C_0}{A_b}$). The point at which the input pump saturates is defined by the turbidostat OD regulation algorithm (which sets a maximum allowable input pump rate). When a pump rate above this point is required OD will thus increase beyond its set-point. Fig. S24a demonstrates cases in which the rate $P_{in}$ required to maintain OD constant rises rapidly (and eventually saturates), as anticipated by (S9).

If we assume $\mu_c$ is constant in time, and given information on pump-rate required to maintain the measured OD constant, (S9) can be used to calculate the biofilm density $\rho_b$ at any point in time. To do this we re-arrange (S9) and multiply by an integrating factor to give:

$$\rho_b(t) = \frac{C_0}{A_b}\int_0^t \Big((P_{in}(p)-\mu_c)\exp\big(-\int_p^t P_{in}(q)-\mu_c+\frac{A_c}{A_b}\mu_s \mathrm{d}q\big)\Big)\mathrm{d}p \tag{S10}$$

which can be evaluated numerically. Alternatively, if we are willing to assume that the biofilm cells undergo exponential growth according to:

$$\dot{\rho}_b = (\mu_b - \mu_s)\rho_b \tag{S11}$$

Combing (S9) and (S11) provides a second approach to calculating $\rho_b$, which gives:

$$\rho_b(t) = \frac{\frac{C_0}{A_b}(P_{in}(t)-\mu_c)}{P_{in}(t)-\mu_c+\mu_b-\mu_s+\frac{A_c}{A_b}\mu_s} \tag{S12}$$

which is easier to implement numerically than (S10).

During an experiment we can estimate $\mu_c$ in the appropriate units by averaging $P_{in}$ at the beginning of the experiment (e.g. first few hours) prior to biofilm formation. The value of $C_0$ can be calculated by combining (S5) and (S6) and again assuming measured OD is being maintained constant to give:

$$C_0 = \mathrm{OD}\cdot\ln(10) \tag{S13}$$

Both (S10) and (S12) have one free parameter which requires estimation. For (S10) we assign $\alpha = \frac{A_c}{A_b}\mu_s$, and for (S12) $\beta = \mu_b - \mu_s + \frac{A_c}{A_b}\mu_s$. A large $\alpha$ implies that shedding of cells by the biofilm ($\mu_s$) is significant (i.e. a large fraction of cells born in the biofilm are shed), which in turn implies that $\alpha \approx \beta$ (since $\mu_b - \mu_s \approx 0$ if shedding is significant). In the large-shedding limit (i.e. $\dot{\rho}_b \to 0$) (S12) becomes an exact solution of (S10). It is worth noting that both $\alpha$ and $\beta$ depend on $\mu_s$ as weighted by the relative optical density of the two cell types ($A_c/A_b$).

In Fig. S24b,c we compare the application of (S10) and (S12) to calculate Biofilm OD based upon the input pump rate (Fig. S24a). We observe that for large $\alpha, \beta$ a lower final Biofilm OD is achieved when pumping saturates; this corresponds to a case where the biofilm is shedding cells into the media at a rate too large for the input pump to compensate. The models with large $\alpha, \beta$ are also perturbed to a lesser degree by small variations in growth/pump rate prior to biofilm onset (i.e. first $\approx 15$ hours of the experiment). Based upon this data alone it is not possible to directly measure either $\alpha$ or $\beta$. That said, at the end of most biofilm experiments (i.e. when pump rate saturates) we observed low densities of cells in solution ($\rho_c$), with OD in the range [0.1,0.2] (note that all experiments were performed with measured OD set by the turbidostat to 0.5), which would thus correspond to a final measured biofilm density in the range [0.3,0.4]. Consequently, we select a mid range $\alpha = 8$ for use in Fig. 3d of the main text. However, it is worth noting that the shedding rate (in the form of $\mu_s$) may change in time as cells adapt to the continuous-culture environment, and will be significantly affected by the rate of stirring (i.e. the shear conditions experienced by the biofilm[17]).

Note that the above analysis assumes both that $\mu_c$ is approximately constant in time and within the OD range of cells considered (i.e. for cell densities less than or equal to the measured OD set-point), and that the OD in question is sufficiently small for the Beer-Lambert relation to (approximately) hold.[13] Ultimately, this analysis is primarily meant as a means for identifying the impact of biofilms on our device's operation (i.e. to alert the user that unwanted biofilms may be forming), rather than a methodology for precise measurement of biofilm formation and growth. As a measurement process it has obvious weaknesses, such as the fact that as the biofilm grows there is a corresponding decrease in the density of cells in the media, which may in turn impact the biofilm growth rate (though in many cases nutrient availability may primarily determine biofilm growth once an initial population is established[18]).

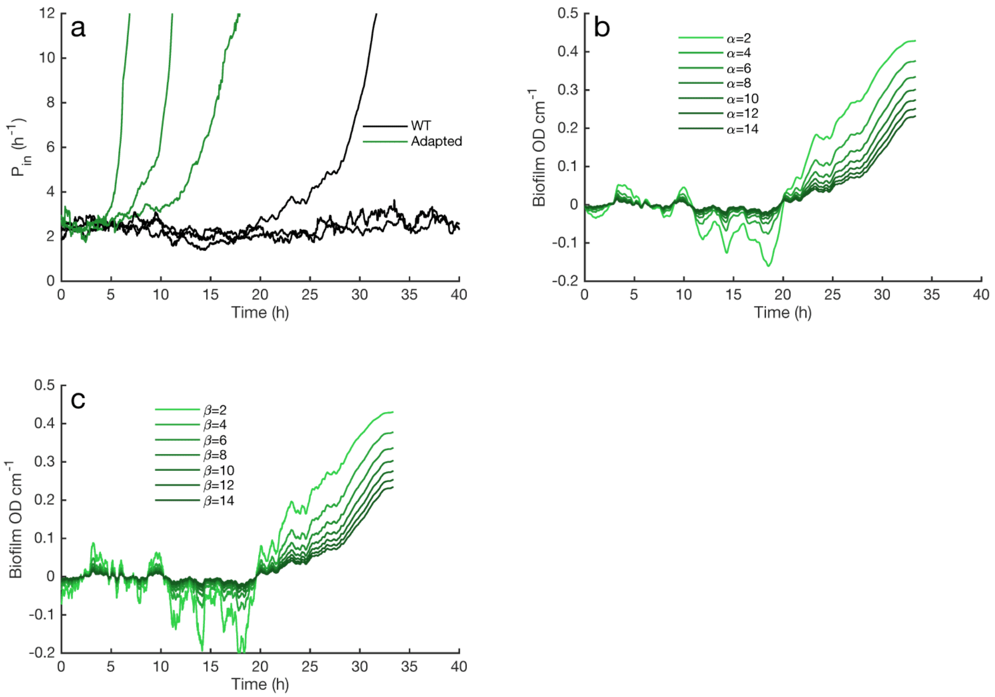

Figure S24: **Biofilm Analysis.** **(a)** Specific pump rate (in units of $h^{-1}$, as in (S8)) for the experiments in Fig. 3d of the main text. **(b)** Biofilm OD calculated using (S10) (the exact solution to (S9)) for various values of $\alpha$. **(c)** Biofilm OD calculated using (S12) (the approximate solution to (S9)) for various values of $\beta$. Note that in all cases pump data has been smoothed using a 1 hour-width moving average filter.

### Note S15 Fluorescence Measurements

##### Measurement Theory

Fluorescence measurement in Chi.Bio is done by activating an appropriate excitation source (one of the coloured LEDs), and then measuring light intensity in two wavelength bands, the *base* band ($I_B$) and the *emission* band ($I_E$). Typically the emission band is set to a wavelength where the protein's fluorescent emission is anticipated, whilst the base band measures the total power of the excitation source (and so usually employs the Clear filter). Normalisation by the absolute excitation intensity means that fluorescence readings are minimally perturbed by changes in culture optical density, or device-to-device variability in excitation source intensity. As an example, GFP measurements typically employ the 457nm LED for excitation, and use the clear filter for the base band measurement and 550nm filter for the emission measurement.

We can (approximately) express the light measured at the baseband as:

$$I_B = I_S \cdot S_B(OD) \tag{S14}$$

where $I_S$ is the absolute intensity of the excitation source (before it enters the test tube), and $S_B$ is a function that describes the quantity of light scattered from the media and cells at the baseband wavelength, which depends on the $OD$ of the culture. In practice $S_B$ may depend on the presence and concentration of a fluorescent protein (with fluorescence emission increasing the total intensity measured by the spectrometer), though we will assume this contribution to be small. The approximate form of $S_B(OD)$ can be observed in Fig. S10a (for the scattering measurements); there is a constant offset due to scattering by the media and test tube, which combines with scattering from the cells (which is $OD$ dependent).

We can express the light measured in the emission band as:

$$I_E = I_S \cdot S_E(OD) + I_S \cdot f_{cell}(OD) + I_S \cdot f_{FP}(OD \cdot FP) \tag{S15}$$

where $S_E$ is a function that describes the quantity of light scattered from the media and cells at the emission wavelength. $f_{cell}$ is a function describing the intensity of fluorescence measured in the excitation band due to cell auto-fluorescence which depends on $OD$. $f_{FP}$ is a function describing the intensity of fluorescence measured in the excitation band due to a fluorescent protein's emission, which depends on the product of $OD$ and the amount of fluorescent protein per cell, $FP$. We expect that both $f_{cell}$ and $f_{FP}$ will be directly proportional to their argument (i.e. $OD$ or $OD \cdot FP$ respectively), without a constant offset (as in $S_B$ and $S_E$). Note that in this analysis we are implicitly assuming that $OD$ is proportional to the density of cells in solution at the concentrations considered.

Typically we express fluorescence per cell as the ratio of these two measured quantities:

$$Fluorescence = \frac{I_E}{I_B} \tag{S16}$$

Combining (S14), (S15), (S16) the power of the excitation source $I_S$ is elimiated, and we are left with:

$$Fluorescence = \frac{S_E(OD) + f_{cell}(OD)}{S_B(OD)} + \frac{f_{FP}(OD \cdot FP)}{S_B(OD)} \tag{S17}$$

where we have sorted the expression into a first term depending only on $OD$, and a second term that additionally depends on the quantity of fluorescent protein. Ideally the expression in (S17) would be a linear function of $FP$ (as we wish to measure fluorescent per cell) and independent of $OD$, which we now investigate.

In Fig. S25a-d we consider a case in which cells start growing from low $OD$ (and are then regulated to an $OD$ of 0.4 cm$^{-1}$), initially without GFP expression. We observe that during initial growth (Fig. S25b) the quantity $I_E/I_B$ is near-constant (Fig. S25d). During this period $I_E$ increases due to light scattered by the cells which is able to pass through the measurement filter ($S_E(OD)$, we anticipate some filter bleed-through from the excitation source to the emission band as illustrated in Fig. S14b). We also plot the quantity $I_E/OD$; this reduces significantly because $S_E$ is not directly proportional to $OD$ in (S15) (due to the baseline scattering from media and test tube).

In Fig. S25e-h we consider a case in which cells (expressing a low-mid level of GFP) start from an $OD$ of 0.4, and are then allowed to grow exponentially into stationary phase (i.e. turbidostat functionality is disabled). Rapid cell growth ensues (Fig. S25f), causing a concomitant increase in $OD$ and accompanying rise in $I_E$ (Fig. S25h). $I_E/I_B$ increases during this period, likely due to the impact of the baseline scattering in $S_B$ (which becomes less significant at high $OD$), though the impact of higher OD is largely mitigated (i.e. $I_E/I_B$ increases $\approx 1.35$ fold in Fig. S25h, whilst $OD$ increases $\approx 4$ fold in Fig. S25f). In this case $I_E/OD$ remains close to constant (though it may not be true that the rate of GFP production *should* be constant across the range of $OD$'s considered). Fluorescence increases significantly as cells enter stationary phase (Fig. S25g) which we hypothesise is due to an increased production rate of the fluorescent protein combined with adequate time available for all fluorophores to mature.

Following the above analysis we elect to quantify fluorescence using (S16) (rather than as $I_E/OD$) for the following reasons; this measurement is far less sensitive to $OD$ changes when fluorescence levels are low (i.e. Fig. S25d), allowing a constant baseline fluorescence for wild-type cells to be accurately subtracted; it is minimally variant across a range of typical $OD$'s (i.e. Fig. S25h); it includes normalisation for inter-device variation (investigated in Fig. S17) in absolute measurement/excitation intensity (via the factor $I_S$, which is not removed when normalising by $OD$). It may be possible to further reduce the deviation of this metric in Fig. S25h by subtracting off the baseline scattering component of both the baseband and emission measurements ($S_B$ and $S_E$), but this would require additional media-specific device calibration.

As demonstrated for the OD laser (Fig. S12c), much of the shot-to-shot variability in fluorescence measurement arises due to inhomogeneities in culture density. The degree of variability depends upon the measurement filters chosen, as well as the absolute level of fluorescence and culture density. This is investigated in Fig. S26, where we assess the inter-shot noise in fluorescence measurements. If multiple emission wavelengths are feasible the choice of which to use is governed by a trade-off between reducing bleed-through of the excitation source into the measurement filter (which is minimised by using higher emission measurement wavelengths), and maximising the fluorescence signal from the protein itself (which for many fluorescent proteins is larger at wavelengths closer to their excitation spectrum, depending on their stokes shift). In the case of GFP we observe a significantly improved signal for larger measurement wavelength (550nm), though the difference in the case of RFP filter choice is minimal. Comparing Fig. S26c,d to e,f highlights another benefit of normalising measurements by the excitation intensity as in (S16); noise due to spatial variation in culture density (which causes a correlated change in both the base band and emission measurement) is decreased.

Following the above, in this study we have taken two steps to post-process fluorescence data (as described in the *Online Methods* of the main text). In the first, we use a time-averaging filter to smooth measurements, which greatly reduces the shot-to-shot variability. Following this a baseline fluorescence value is subtracted (corresponding to the left term in (S17)) to remove the effect of media and cell auto-fluorescence,[20] as well as filter bleed-through (excitation light that passes through the emission measurement filter), which (as demonstrated in Fig. S15) varies minimally between devices. The effect of these two post-processing steps can be observed by comparing Fig. 3e-h of the main text (processed data) with Fig. S27 (raw data).

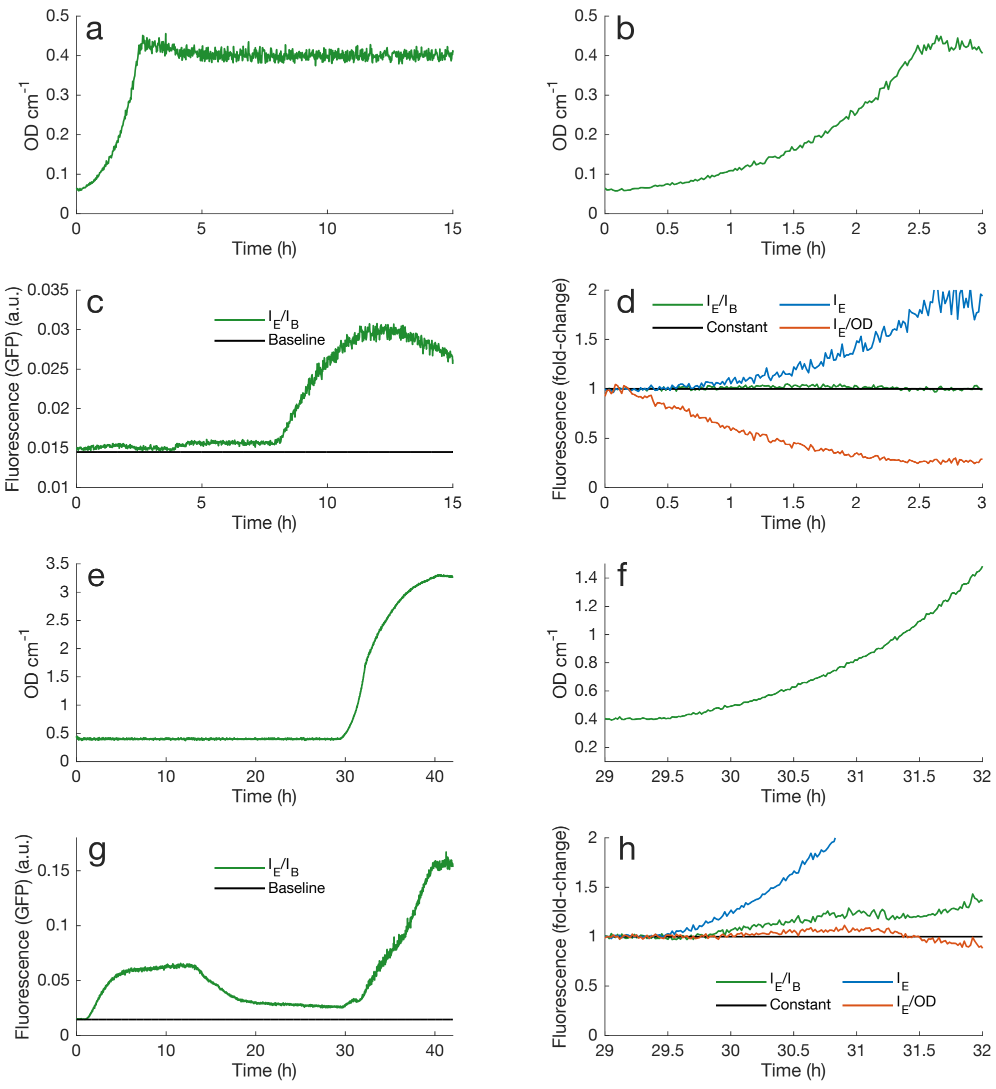

Figure S25: **Fluorescence Measurement Technique.** All data is for *E. coli* expressing GFP (excited at 457nm, with Clear baseband and 550nm emission band) grown in EZ media. **(a)** An experiment starting at low OD, for which the first three hours are re-plot in **(b)**. **(c)** Measured GFP throughout the experiment, with Baseline indicating that device's average measurement for wild-type cells. GFP is chemically induced at $t \approx 8$ hours. Note that fluorescence increases slightly at $t \approx 4$ hours due to addition of aTc (i.e. as in Fig. 3k). **(d)** Comparison of different standardisation methods for fluorescence measurements over the first three hours of the experiment in **c**, each measurement is divided by its value at $t = 0$ to give a fold-change metric. **(e)** A different experiment, in which turbidostat regulation is disabled at $t = 29.5$ hours at an intermediate GFP level. **(f)** Detail of the period around turbidostat de-activation. **(g)** Fluorescence for the experiment in **e**, in which GFP is induced at $t \approx 1$ hour and RFP is induced at $t \approx 13$ hours, leading to a reduction in GFP (as in Fig. 3j). **(h)** Similar fold-change analysis to **d**, demonstrating that $I_E$ tracks the OD profile in **f**, whilst $I_E/I_B$ remains approximately constant.

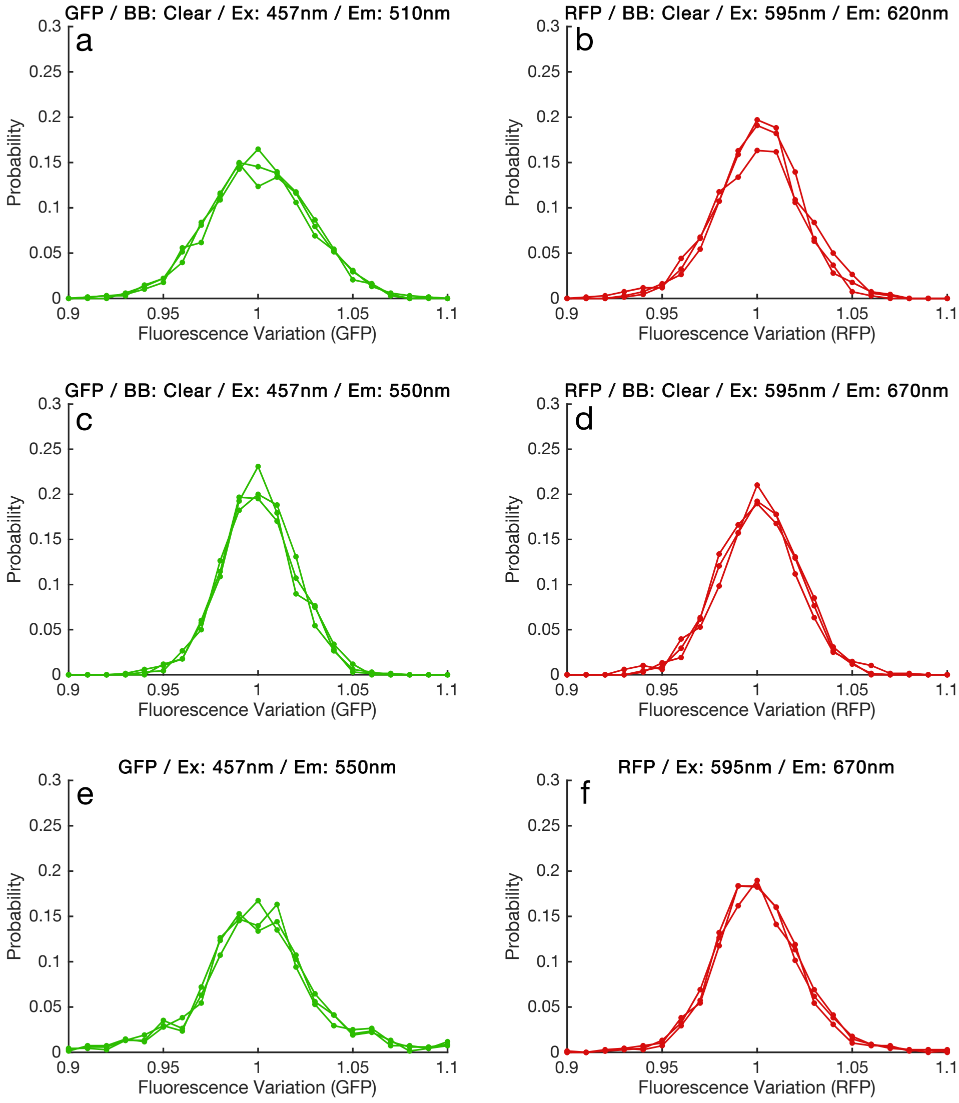

Figure S26: **Fluorescence Measurement Analysis.** Shot-to-shot variability of fluorescence measurements for three devices (the data in Fig. S28). Fluorescence Variation is calculated by dividing the raw experimental data by the same data with a 20 minute-width smoothing filter applied. **(a)** GFP measurement with excitation by 457nm LED, Clear filter baseband, and 510nm emission filter. **(b)** RFP measurement with excitation by 595nm LED, Clear filter baseband, and 620nm emission filter. **(c)** GFP measurement with excitation by 457nm LED, Clear filter baseband, and 550nm emission filter. **(b)** RFP measurement with excitation by 595nm LED, Clear filter baseband, and 670nm emission filter. **(e,f)** Same as **c,d**, but without the base-band normalisation of measurements. Noise increases (particularly in the GFP case) due to inhomogeneities in culture density.

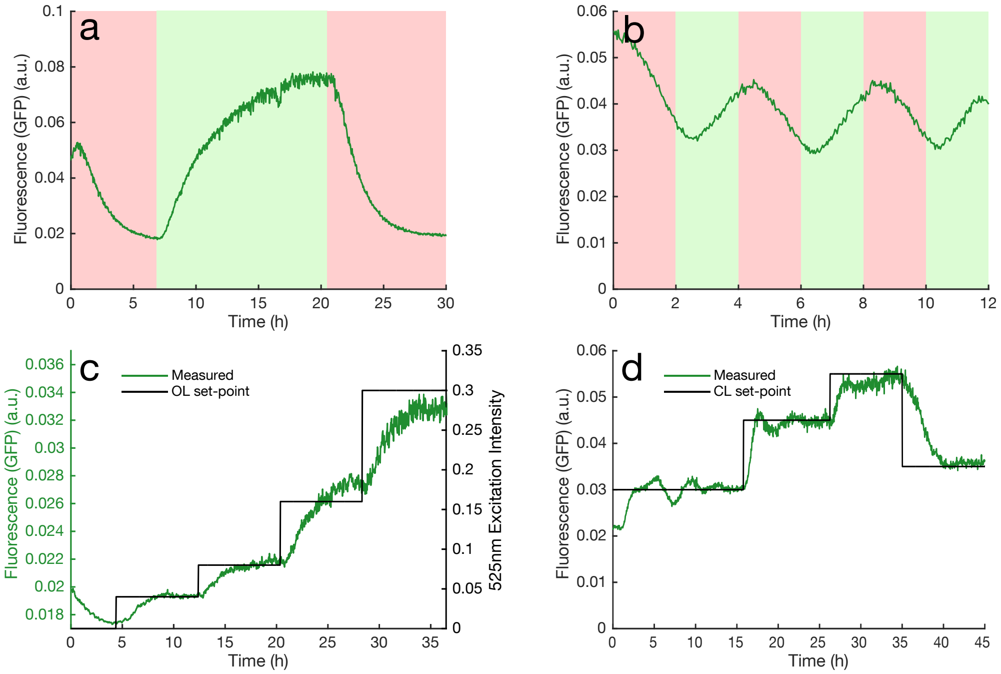

Figure S27: **Application 2 - Raw Data.** Raw data (directly as collected by Chi.Bio) for Application 3 in the main text. In this case no smoothing filter has been applied, nor has the baseline fluorescence level (≈ 0.017) been subtracted.

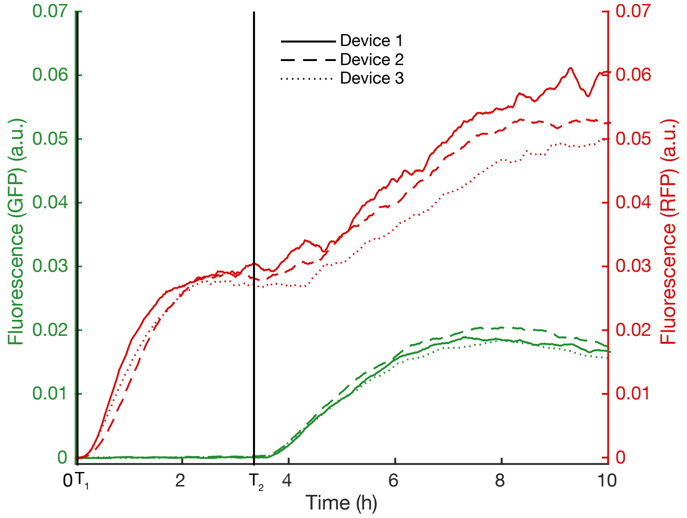

Figure S28: **Application 3 - Repeat.** Triplicate repeat of experiment similar to Application 3, in which aTc is added at $T_1$ and L-Rhamnose added at $T_2$. In all cases we observe a similar increase in RFP following GFP addition, which we hypothesise is due to a reduced concentration of the repressing transcription factor TetR.

#### Note S16 Other Applications

In addition to the applications outlined herein, Chi.Bio has the potential to be used for a broad range of studies across the biological sciences. Table S1 outlines a selection of potential applications, and describes simple modifications that could be made to the platform to enable others.

| Application | Notes |
|---|---|
| Single-cell analysis | If single-cell measurements are required one of Chi.Bio's additional pumps can be employed to direct samples to a measurement apparatus at regular intervals (e.g. as done by Milias-Argeitis *et al*[10]). Alternatively, mechanisms for measurement of intra-population variability using only bulk measurements are also under development.[21] |
| Inter-chamber mixing | Similar to the above, pumps 3 and 4 can be routed between chambers to (for example) automate passaging for biofilm prevention (as implemented by the eVOLVER platform[8]). |
| Chemostat operation | The OD control algorithm can easily be re-programmed to provide constant input/output pump rates to provide a chemostat setup. |
| Biosynthesis Optimisation | Optimisation of biosynthesis processes often requires analysing the impact of multiple experimental parameters, such as temperature, culture agitation, or nutrient combinations. Chi.Bio can be programmed to scan through a parameter space (e.g. using two pumps to vary the properties of its input media) whilst recording growth rate and spectral data. |
| Algae/Cyanobacteria | Chi.Bio can be used as-is to grow photosynthetic organisms, and provides an unparalleled ability to tune its light source's spectrum (thereby controlling growth[22]). Due to the absorbance of photosynthetic organisms, infrared wavelengths are regularly used to measure their optical density. If this is required Chi.Bio can easily be retrofit by replacing its internal laser diode (which is a standard TO-18 5.6mm package) with an infrared version (e.g. the 780nm 5mW diode produced by Rohm) that can be measured by the spectrometer (which is sensitive up to $\approx 1\mu$m). |
| Spectral Analysis | Measurement of absorption spectra can be used to infer many features of a cell culture, such as average cell size and contents,[23–25] ratios of constituent species,[26] or pH when specialised dyes are added.[27] Integration of reactive sensor patches into the growth chamber could provide an additional means of sensing of pH and dissolved Oxygen.[28] Chi.Bio can be used as-is for many such applications, so long as the combination of measurement filters and light sources is sufficient. |
| Anaerobic / gas mixing | Chi.Bio uses standard laboratory test tubes with the common 22-400 thread. Consequently there are many commercial lids that can be substituted for our custom-made lid, if liquid inputs are not desired. Developing an air-tight pumping/closure apparatus (to allow precise control of atmosphere within the growth chamber) is a direction for future development of the platform. |
| Experimental Evolution | Many continuous-culture experimental evolution experiments can be implemented in Chi.Bio as-is by using its existing control hardware/software to vary parameters or chemical concentrations.[29, 30] Procedures which require a higher degree of sterility, or closed atmosphere, would require development of an air-tight pumping/closure apparatus as above. |
| Education | Chi.Bio is very easy to set up and use, making it suitable for teaching at graduate, undergraduate, and late high school levels. The reactor's core features (heating, stirring, spectrometry) can be used as a stand-alone low-cost measurement/manipulation platform, which allows students to clearly visualise how measurements are taken (e.g. they can remove the lid and observe fluorescence by eye). |

Table S1: **Additional applications.**

### Note S17 Strains Used In This Study

| Strain | Description | Reference |
|---|---|---|
| *E. coli* MG1655 | Wild-type K12 *E. coli*, used for growth/biofilm experiments. | Blattner *et al.*[31] |
| *E. coli* BL21(DE3) | Used for production of fluorescent proteins. *fhuA2 [lon] ompT gal (λ DE3) [dcm] ΔhsdS λ DE3 = λ sBamHIo ΔEcoRI-B int::(lacI::PlacUV5::T7 gene1) i21 Δnin5* | New England Biolabs |
| *E. coli* BW29655 | Used for optogenetics experiments. *Δ(araD-araB)567 ΔlacZ4787(::rrnB-3) λ- Δ(envZ-ompR)520(::FRT) Δ(rhaD-rhaB)568 hsdR514* | CGSC# 7934, Zhou *et al.*[32] |

Table S2: **List of strains used in this study.**

### Note S18 Plasmids Used In This Study

| Plasmid | Description | Reference |
|---|---|---|
| pUC19 | $Amp^R$, pMB1, *lacZα* | New England Biolabs |
| pCK302 | $Amp^R$, pBR322, *rhaS*, $P_{rhaBAD}$-*sfgfp* | [33] Shared by Dr C. Kelly |
| pBbA2c_*rfp* | $Cm^R$, p15A ori, *tetR*, $P_{tet}$-*mrfp1* | [34] Shared by Prof J. Keasling |
| pSR58.6 | Used for optogenetics experiments. $Cm^R$, ColE1, $P_{J23100}$-*ccaR*, $P_{cpcG2-172}$-*sfgfp* | [35] Shared by Prof J. Tabor (Addgene plasmid # 63176; http://n2t.net/addgene:63176 ; RRID:Addgene_ 63176). |
| pNO286-3 | Used for optogenetics experiments. $Spec^R$, p15A, $P_{J23106}$-*mini-ccaS*, $P_{J23108}$-*ho1-pcyA* | [35] Shared by Prof J. Tabor (Addgene plasmid # 107746; http://n2t.net/addgene:107746 ; RRID:Addgene_ 107746). |

Table S3: **List of plasmids used in this study.**

### Supplementary References

[1] Melexis, "Digital plug & play infrared thermometer in a TO-can," 2019.

[2] AMS, "AS7341 11-Channel Spectral Color Sensor," 2019.

[3] LEDEngin, "LZ7 LED Emitter," 2019.

[4] K. Nagasawa, J. Iwase, D. Comparini, and T. Kawano, "Empirical and simulative evaluations of white fluorescence-type light emitting diodes as algal growing light sources based on the photosynthetic oxygen evolution by synechocystis spp. PCC6803," *Environmental Control in Biology*, vol. 53, no. 3, pp. 169–173, 2015.

[5] RayVio, "XR Series UV Emitter," 2019.

[6] N. Vermeulen, W. J. Keeler, K. Nandakumar, and K. T. Leung, "The bactericidal effect of ultraviolet and visible light on Escherichia coli," *Biotechnology and Bioengineering*, vol. 99, no. 3, pp. 550–556, 2008.

[7] F. Cockerill, *Methods for Dilution Antimicrobial Susceptibility Tests for Bacteria That Grow Aerobically; Approved Standard.* CLSI, 9th ed., 2012.

[14] M. Grinberg, *Flask web development: developing web applications with python.* O'Reilly Media, Inc, 2018.

[15] Various, "Gunicorn Project," 2019.
